# Temporal coordination between chromosome mobility and homologous recombination

**DOI:** 10.1101/2022.03.24.485580

**Authors:** Fraulin Joseph, So Jung Lee, Eric Edward Bryant, Robert J. D. Reid, Ivana Sunjevaric, Rodney Rothstein

**Affiliations:** Columbia University Irving Medical Center, Department of Genetics & Development New York, NY 10032, USA; Columbia University, Department of Biological Sciences New York, NY 10027, USA; Columbia University Irving Medical Center, Department of Systems Biology

## Abstract

Homologous recombination (HR), a principal cellular pathway for double-strand break (DSB) repair, is linked to changes in chromosome movement. Although increased chromosome mobility in response to a DSB has been observed in a variety of species, its precise role in HR remains controversial. Here, we find that end resection, the recruitment of recombination proteins, increased chromosome mobility, the pairing of homologs and gene conversion are temporally linked in response to a DSB. In *mre11Δ* mutant cells, which exhibit a delay in the initial processing of a DSB, chromosome mobility and all subsequent recombination events are also delayed. Overexpression of the Dna2 nuclease suppresses the *mre11Δ* delay in end resection and restores the original timing of chromosome mobility and all subsequent downstream HR events. Thus, changing the timing of chromosome mobility results in a corresponding change in essential downstream HR events, reinforcing its mechanistic role in the DNA repair process.

## Introduction

Homologous recombination (HR) is an important pathway for the repair of double-strand breaks (DSBs) that occur as a result of exogenous or endogenous DNA damage (Ciccia & Elledge, 2010). Although many of the steps of HR are well characterized (Lisby & Rothstein, 2015; Wright, Shah, & Heyer, 2018), how a broken chromosome rapidly and efficiently finds its homolog residing microns away in the nucleus poses a fascinating problem. While increased movement of the broken chromosome within the nucleus has been initially proposed to be essential for homology search (Mine-Hattab & Rothstein, 2013; Seeber & Gasser, 2017; Smith & Rothstein, 2017), currently, that view is controversial. Studies in mammalian cells reveal conflicting results, with some studies exhibiting large-scale movements of DSBs and others showing breaks to be mostly immobile (Dion & Gasser, 2013; Merigliano & Chiolo, 2021; Mine-Hattab & Chiolo, 2020). Some studies in yeast report that local increased chromosome mobility at a break site is not required for HR. Changes in the centromeric or telomeric attachments of chromosomes have been proposed to drive chromosome movement suggesting that increased chromosome mobility is not directly connected to HR (Lawrimore et al., 2017; Strecker et al., 2016). Similarly, global reduction in nucleosome density, rather than a local increase in DSB movement has been proposed to be the rate limiting step for homology directed repair (Challa et al., 2021; Cheblal et al., 2020). Other studies report that chromosome positioning within the nucleus is the most important factor for determining the success of HR repair (C. S. Lee et al., 2016). Notably, these yeast studies were performed in haploid cells and only examined ectopic recombination at one time point. Thus, although there are data that support a role for chromosome mobility in the HR pathway, the precise relationship between mobility and recombination is not clear.

To directly test whether mobility is important for HR, one would ideally examine HR in a mutant that specifically blocks mobility. Because such a mutant has not been identified, we took an alternative approach and studied the coordination and timing of key HR events after the induction of a site-specific DSB. We find a tight temporal correlation between the recruitment of recombination proteins, increased mobility, the physical pairing of homologous loci, and gene conversion. Importantly, the timing between these events is preserved even when mobility is delayed in the resection defective mutant, *mre11Δ*. Furthermore, overexpression of Dna2, a nuclease involved in DSB end resection, suppresses the delay in chromosome mobility and simultaneously restores the original timing of all the downstream events without altering their temporal relationship. These results support the hypothesis that increased chromosome mobility promotes the pairing of homologous loci and is dependent on the initial timing of DNA end processing.

## Results

### Chromosome mobility is delayed in *mre11Δ* cells

To explore whether chromosome mobility is mechanistically linked to HR, we tested how cells respond to DSBs when an early step in HR is disrupted. To this end, we examined chromosome mobility in the absence of the Mre11-Rad50-Xrs2 (MRX) complex, one of the first responders to a DSB (Lisby, Barlow, Burgess, & Rothstein, 2004; Lisby & Rothstein, 2009). This complex plays a crucial role in DSB processing for homology-directed repair (Lamarche, Orazio, & Weitzman, 2010) and in the absence of the *MRE11* gene, the MRX complex does not form (Usui et al., 1998). To monitor chromosome mobility, we developed a system to track the movements of a broken chromosome during the repair of a DSB (Mine-Hattab & Rothstein, 2012). Briefly, this system includes a chromosome tag located adjacent to an inducible I-*Sce*I cut site and a tagged Rad52 protein to mark DSBs that are undergoing repair (Fig. 1A). Early S-phase cells were selected that had either no visible Rad52 focus before treatment (Fig. 1B, uninduced, top panels) or a Rad52 focus colocalizing with the tetO array after DSB induction (Fig. 1B, DSB, bottom panels). The tetO array was then tracked in three dimensions, its displacement was measured every 20 seconds and the mean squared displacement (MSD) was plotted for the time intervals from 20 to 900 seconds (Fig. 1C). In the absence of DSBs, WT and *mre11Δ* cells behave similarly and display confined chromosome mobility with a radius of confinement of 500 nm (Fig. 1C, uninduced). In contrast, after 3 hours of DSB induction, WT cells exhibit increased chromosome mobility, while *mre11Δ* cells do not (Fig. 1C, 3 hr post-DSB induction). However, increased mobility is observed in *mre11Δ* cells 1 hour later (Fig. 1C, 4 hr post-DSB induction). Thus, loss of the MRX complex delays but does not completely abolish mobility, showing that this complex is important, but not essential, for increased chromosome mobility.

**Figure 1.**
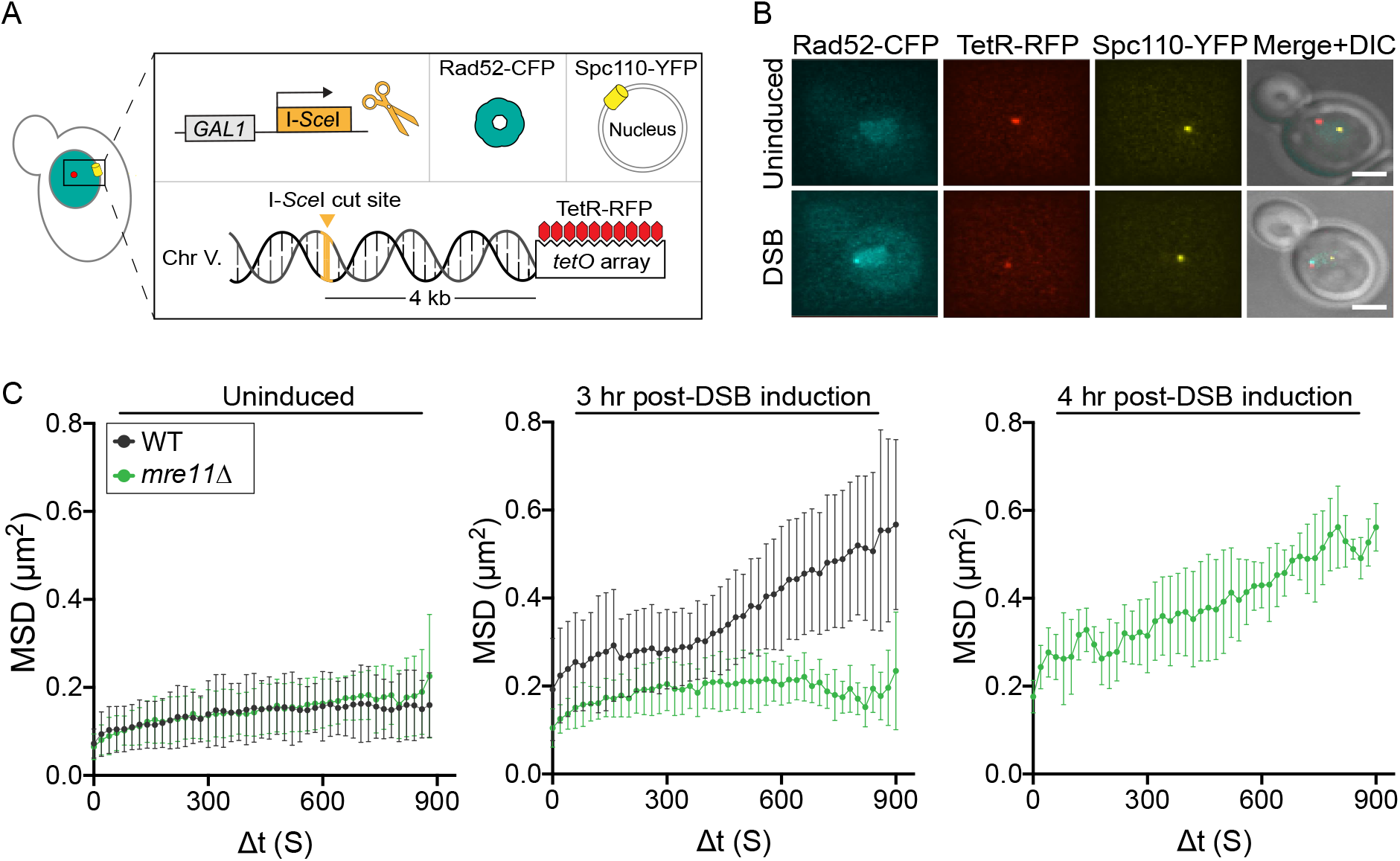
Chromosome mobility is delayed in *mre11Δ* cells. **(A)** Schematic of the strain used to examine chromosome mobility in diploid yeast cells. A multiple tandem array of the Tet-Operator (*tetO*) is integrated adjacent to the *URA3* locus on chromosome V. Constitutively expressed Tet-Repressor proteins fused to RFP (TetR-RFP) bind the *tetO* array (red) allowing its visualization for time lapse imaging by fluorescence microscopy. An I-*Sce*I cut site (Goldstein & McCusker) is located 4 kb from the *tetO* array. To produce a site-specific DNA break, the I-*Sce*I endonuclease is expressed after exposure to galactose. In addition, the strain contains a Rad52-CFP fusion protein (cyan), which accumulates at the I-*Sce*I cut site and acts as an indicator of DSB formation. Spc110 (yellow), a structural component of the spindle pole body (Yoder, Pearson, Bloom, & Davis, 2003), is tagged with YFP and is used as a point of reference when tracking the movement of the *tetO* array. **(B)** Images show deconvolved z-series projections for representative examples of an uninduced cell (no DSB, top) where Rad52-CFP forms a diffuse haze within the cell nucleus and an I-*Sce*I-induced cell (bottom) containing a Rad52 focus that colocalizes with the tetO array. Each fluorescently tagged protein is indicated above the image (Scale bar = 2 microns). **(C)** MSD plots for the *tetO* array in WT (W11577) cells in black or *mre11Δ* (W11578) in green. Cells were untreated (left) (n = 16 for WT and n = 18 for *mre11Δ*), induced for 3 hours (middle) (n = 13 for WT and n = 15 for *mre11Δ*), or induced for 4 hours of I-*Sce*I expression (n = 11 for *mre11Δ*). Error bars represent the 95% CI.

### DNA end-resection is delayed in *mre11Δ* cells

We hypothesized that the delay in chromosome mobility observed in diploid *mre11Δ* cells is the result of a defect in DNA end resection. In haploid cells, DNA end resection is slower and less efficient in the absence of the MRX complex, but can still occur through the activity of the nucleases Dna2-Sgs1 and/or Exo1 (Cejka, 2015; Ivanov, Sugawara, White, Fabre, & Haber, 1994; Zhu, Chung, Shim, Lee, & Ira, 2008). To examine DNA end resection in *mre11Δ* cells, we used a qPCR assay, initially developed in haploids (Gnügge, Oh, & Symington, 2018; Zierhut & Diffley, 2008), to measure single-stranded DNA (ssDNA) at two *Dra*I sites located either 0.3 kb or 4.0 kb away from the I-*Sce*I cut site (Fig. S1A). Using this assay in one of the haploid parents of our diploid strain, we find that resection is delayed in *mre11Δ* cells (Fig. S1C, D) similar to that observed in haploid *mre11Δ* cells after an HO cut at the mating type locus (S. E. Lee, Bressan, Petrini, & Haber, 2002; Moreau, Morgan, & Symington, 2001). To examine end processing in a diploid, we designed primers that only amplify the I-*Sce*I recognition sequence found exclusively on the *tetO* array-containing chromosome and not the homologous *lacO* chromosome (Fig. 2A, cut site, yellow). As observed in haploid cells (Fig. S1B), the percent cutting of the I-*Sce*I site was the same in both WT and *mre11Δ* diploids (Fig. 2B). In WT diploids, the percent ssDNA at both the 0.3 kb and 4.0 kb *Dra*I sites increases, peaks at 2 and 3 hours of DSB induction, respectively, and then declines (Fig. 2C, D, black line).

**Figure 2.**
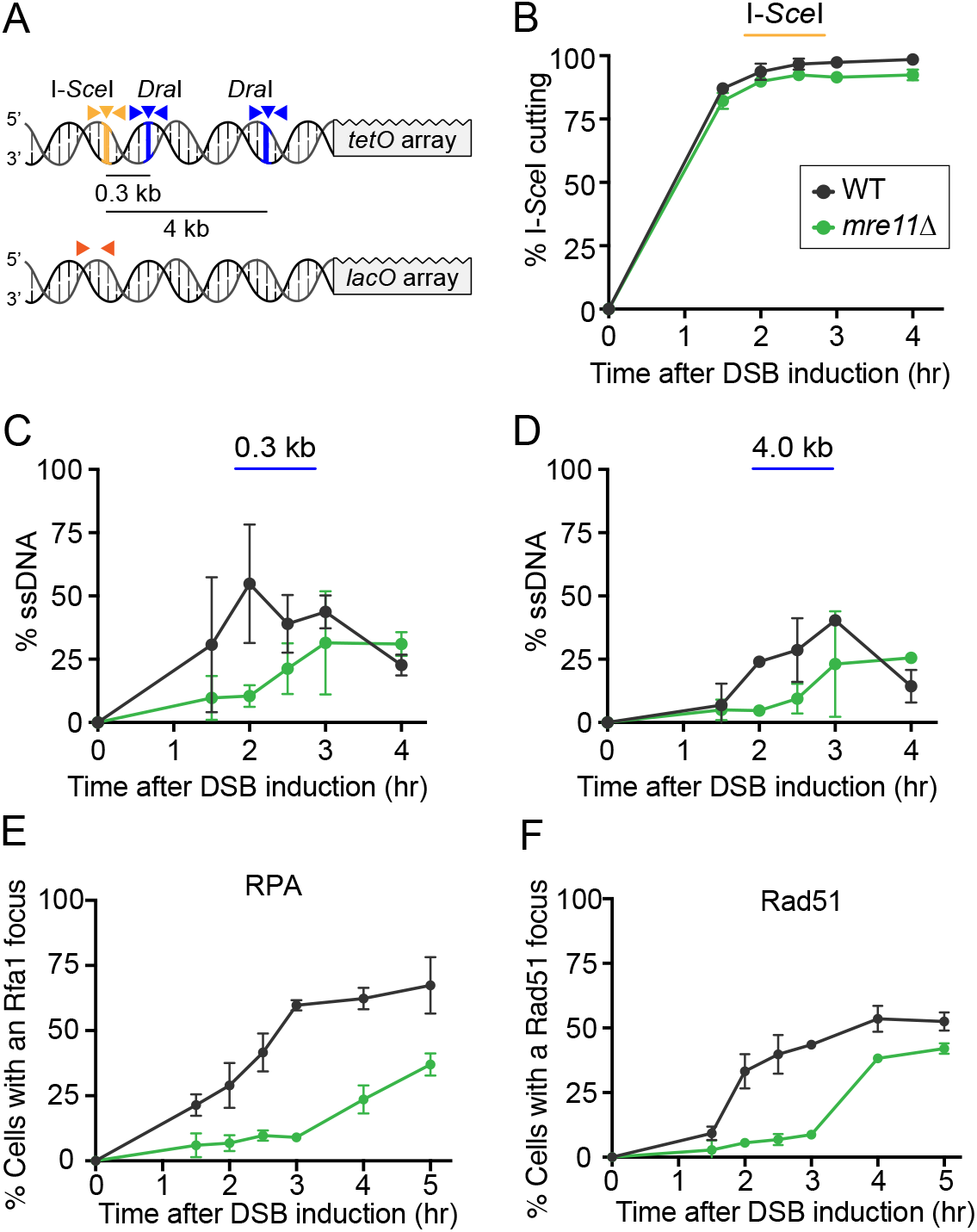
DNA end-resection is delayed in *mre11Δ* cells. **(A)** Schematic of the diploid strain used to measure ssDNA via qPCR. *Dra*I sites (blue lines) are located either 0.3 kb or 4.0 kb away from the I-SceI cut site (yellow line). PCR primers that flank each *Dra*I site (blue arrowheads) are used for amplification before and after *in vitro* digestion with *Dra*I. The percent of *Dra*I-resistant DNA reveals the amount of DNA resection at that site. Amplification of DNA from primers that flank the I-*Sce*I site (yellow arrowheads) is used to measure I-SceI cutting. To measure I-*Sce*I repair, primers were designed that only amplify the donor sequence present on the homologous chromosome (orange arrowheads). **(B)** Percentage of I-*Sce*I cutting plotted vs. time after DSB induction in WT (W11579) and *mre11Δ* (W11580) cells. **(C-D)** Percentage of ssDNA present at the 0.3 kb and 4.0 kb *Dra*I sites plotted vs. time in the same cells from (B). Error bars represent the standard deviation from 3 biological replicates. **(E-F)** The percent of cells with an Rfa1-YFP (E) or Rad51-YFP (F) focus colocalizing with the I-*Sce*I cut site after DSB induction are plotted. WT (W11581, W11582) cells are shown in black and *mre11Δ* (W11583, W11584) in green. At each time point, at least 100 cells were analyzed for each strain. Error bars represent the standard deviation from 3 biological replicates.

We interpret the decline of ssDNA in WT diploid cells as repair of the I-*Sce*I cut site using the homolog since this decline is not observed in haploids, where no repair template is present (Fig. S1C, D). For diploid *mre11Δ* cells, the percent increase in ssDNA at both *Dra*I sites is delayed approximately one hour compared to WT (Fig. 2C, D, green lines), similar to the delay observed for increased mobility (Fig. 1C). Thus, these results demonstrate that the delay in mobility correlates with the timing of DNA end resection in *mre11Δ* cells.

### Rad51 and Rfa1 protein recruitment to a DSB site are delayed in *mre11Δ* cells

DNA end resection at a DSB produces ssDNA regions that are rapidly bound by repair proteins. Since resection is delayed in an *mre11Δ* strain, we predicted that accumulation of ssDNA binding proteins at DNA ends would be similarly delayed. We first looked at Rfa1, a subunit of the RPA complex, binds to ssDNA (Brill & Stillman, 1989) and provides a platform for the recruitment and activation of the checkpoint kinase Mec1 (Zou & Elledge, 2003). This recruitment activates the DNA damage response, which is necessary and sufficient to induce chromosome mobility (Dion, Kalck, Horigome, Towbin, & Gasser, 2012; Seeber, Dion, & Gasser, 2013; Smith, Bryant, & Rothstein, 2018). We also examined Rad51, which forms filaments on ssDNA following DNA end resection (Laurini et al., 2020; Symington, 2016; Symington, Rothstein, & Lisby, 2014) and is essential for homology search as well as chromosome mobility in yeast (Dion et al., 2012; Mine-Hattab & Rothstein, 2012). We find that the accumulation of Rfa1 and Rad51 foci at an I-*Sce*I induced break site is delayed in *mre11Δ* cells compared to WT (Fig. 2E, F). Thus, in the absence of the MRX complex, Rfa1 and Rad51 protein accumulation at the DSB site is delayed, paralleling the delay in ssDNA formation and chromosome mobility. Furthermore, these results are consistent with the idea that Rfa1 and Rad51 loading on resected DNA is an important precursor to chromosome mobility.

### Homolog pairing is delayed in *mre11Δ* cells

We hypothesized that chromosome mobility affects damage-induced pairing between homologs and predicted a corresponding delay in pairing in *mre11Δ* cells. To test this idea, we employed a system to visualize real-time homolog pairing in living diploid yeast cells (Mine-Hattab & Rothstein, 2012) (Fig. 3A, B). In WT cells, after induction of a I-*Sce*I DSB, the percentage of paired homologs increases sharply after 90 minutes, peaking at 2.5 hours and subsequently decreases. This behavior likely reflects the physical pairing of the homologous loci to initiate repair and their subsequent separation after the process is complete (Fig. 3C, black line).

**Figure 3.**
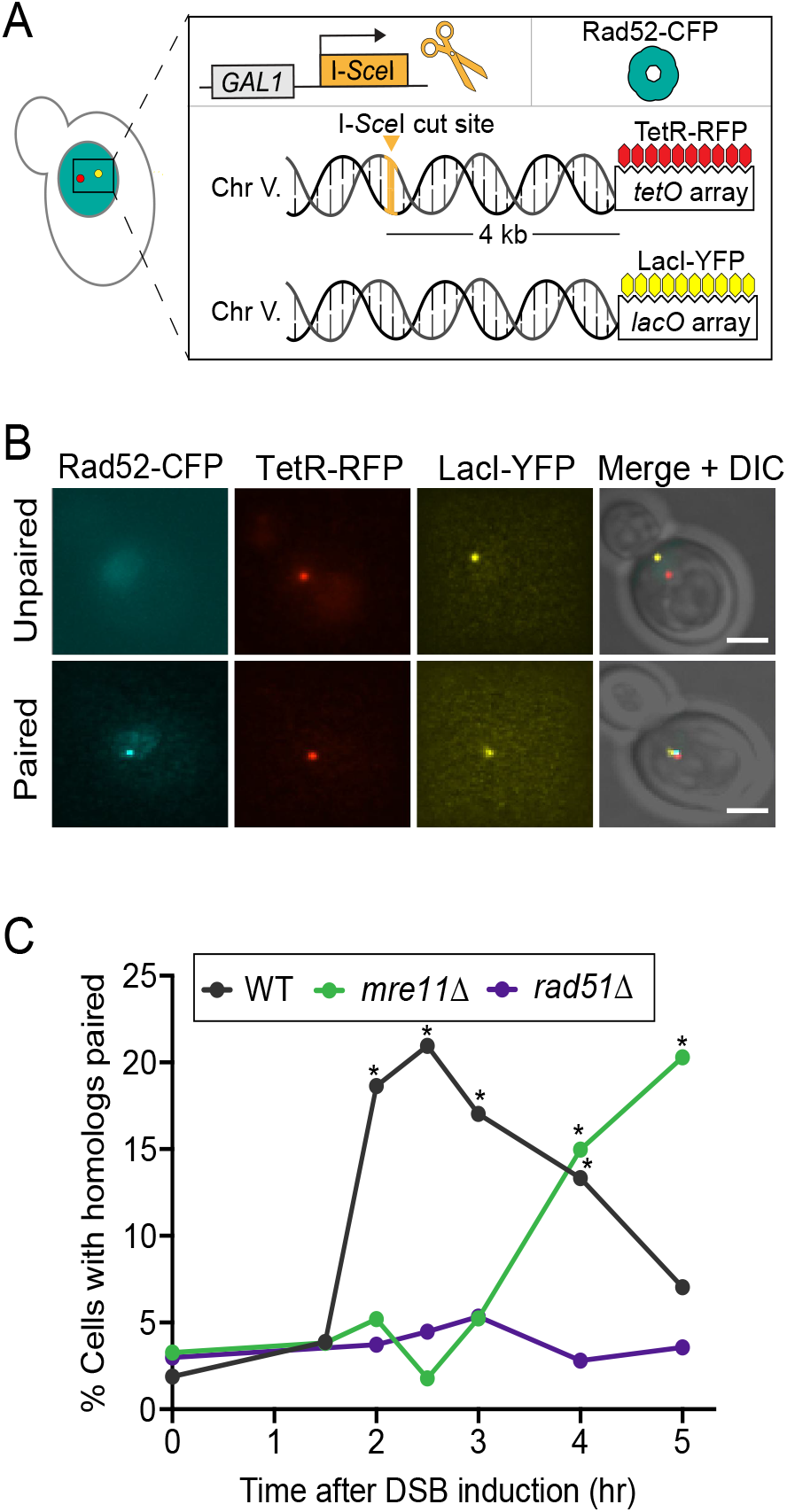
Homolog pairing is delayed in *mre11Δ* cells. **(A)** Schematic of the strain used to examine the pairing behavior of homologs. A multiple tandem array of the Lac-operator (*lacO*) is inserted on the chromosome V homolog at the same position as the *tetO* array in the strain described in Figure 1. The two homologous loci are visualized by the binding of Tet and Lac repressor proteins fused to fluorophores (TetR-RFP and YFP-LacI, respectively). Homolog pairing is revealed by the co-localization of the red and yellow foci. **(B)** Images show deconvolved z-series projections for representative examples of cells with homologs unpaired (top) and paired after an I- *Sce*I-induced DSB (bottom). Each fluorescently-tagged protein is indicated above the image. The merged CFP, RFP and YFP images are shown overlaying the DIC image (Scale bar = 2 microns). **(C)** The percent pairing of the homologs after DSB induction is plotted. WT (W11579) cells are shown in black, *mre11Δ* (W11580) in green, and *rad51Δ* (W11585) in purple. At each time point, at least 100 cells were analyzed for each strain. The threshold for homolog pairing was determined as described in Figure S2. Fisher’s exact proportion test was used to determine the statistical significance. All data points marked with an asterisk show an increase in the percent pairing that is significantly greater than the < 5% background level of colocalization seen in the *rad51Δ* negative control and have a p-value less than 0.05.

Pairing is dependent on the Rad51 recombinase since *rad51Δ* cells do not show increased homolog pairing (i.e., colocalization) after DSB induction, remaining below the background throughout the time course (Fig. 3C, purple line). In *mre11Δ* cells, pairing was not observed until 4 hours after DSB induction (Fig. 3C, green line). Notably, this delay in pairing parallels what we observed for end processing, the recruitment of repair factors, and increased chromosome mobility at the break site observed in *mre11Δ* cells (Fig. 1C and Fig. 2C-F). These data are consistent with the idea that mobility is temporally linked to critical events in HR, including homolog pairing.

### DSB repair via gene conversion is also delayed in *mre11Δ* cells

Finally, we tested whether the genetic recombination event itself is temporally linked to mobility. In our diploid strains, genetic recombination is measured by gene conversion, which is detected by the loss of the I-*Sce*I cut site on the *tetO*-containing chromosome as a result of DSB repair from the *lacO*-containing chromosome (Mine-Hattab & Rothstein, 2012) (Fig. 4A). This process is dependent on the Rad51 recombinase and independent of NHEJ (Fig. S3). In both WT and *mre11Δ* cells, genomic blots reveal a 7 kb band, which indicates *in vivo* cutting of the I-*Sce*I site (Fig. 4B, left panels). Quantitation of the blots shows that I-*Sce*I cutting for WT and *mre11Δ* is the same during the first hour of DSB induction (Fig.4C, left panels). In WT cells, the percent cut DNA begins to decrease after 1.5 hours due to loss of the I-*Sce*I site via gene conversion from the homolog, which lacks the cut site. In *mre11Δ* cells, the percent cut DNA begins to decrease after 3 hours of DSB induction (Fig. 4C, left panels). To verify that the decrease in I- *Sce*I cutting in both WT and *mre11Δ* was indeed due to gene conversion, the same samples from the left panel of Figure 4B were digested with purified I-*Sce*I enzyme *in vitro* (Fig. 4B, right panels). In WT cells, the 8 kb I-*Sce*I-resistant band due to gene conversion (labeled GC) is detected after 2 hours of DSB induction, whereas in *mre11Δ* cells that same band appears 1.5 to 2 hours later (Fig. 4B, C, right panels). Thus, in *mre11Δ* cells, the delay in gene conversion corresponds with the delay observed for increased chromosome mobility and homolog pairing, further supporting the notion that all of these processes are linked (Fig. 1C and Fig. 3C).

**Figure 4.**
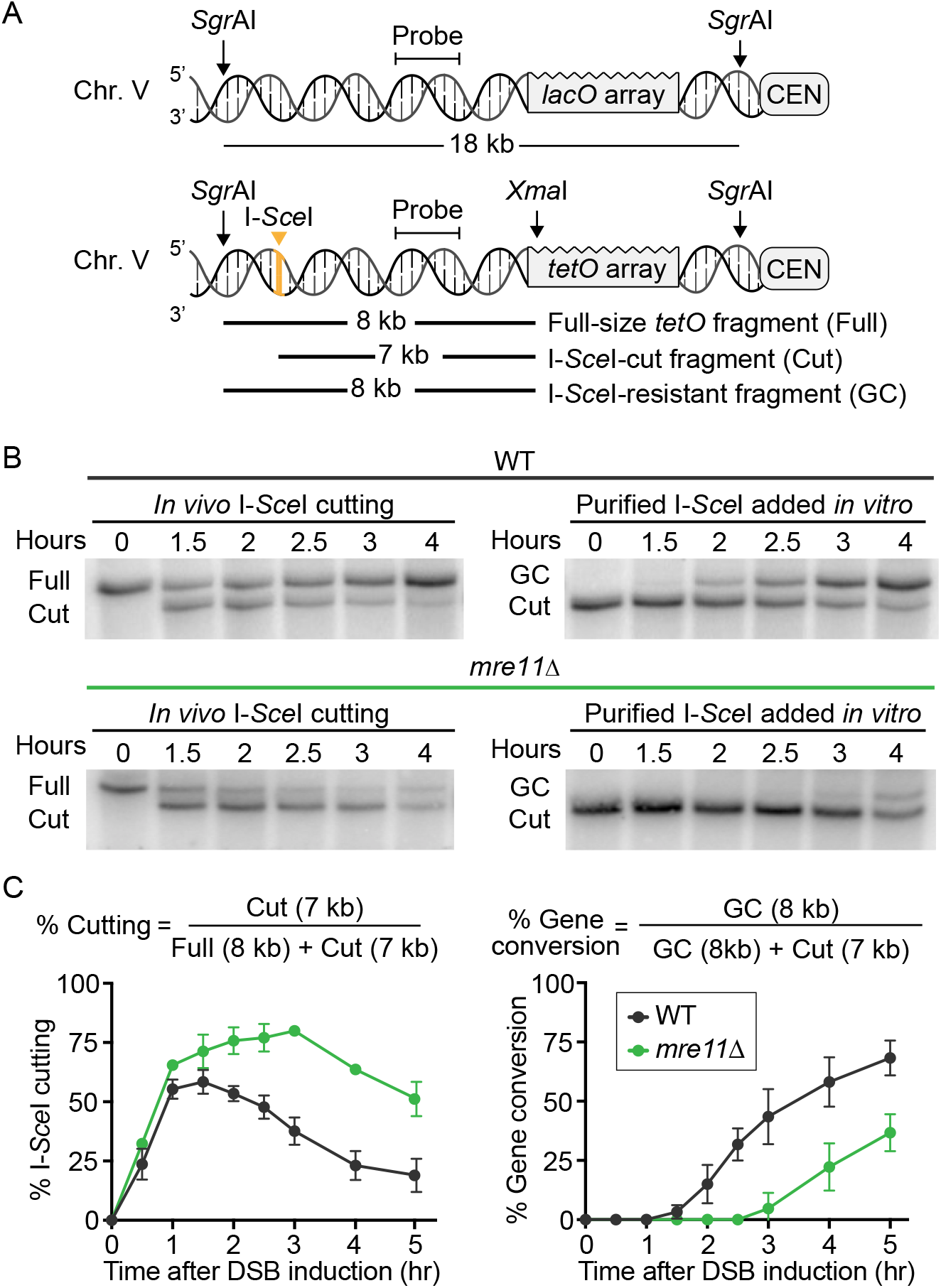
DSB repair via gene conversion is delayed in *mre11Δ* cells. **(A)** Schematic (not to scale) of chromosome V homologous loci containing repressor-binding arrays, in which one homolog has an I- *Sce*I cut site. The location of the *Sgr*AI and *Xma*I restriction sites, DNA probe, and the expected fragment sizes after digestion are also shown. *In vivo* cleavage by I-*Sce*I produces a 7 kb DNA fragment after digestion with *Sgr*AI and *Xma*I. However, if the I-*Sce*I site is repaired using the homologous *lacO* chromosome, an 8 kb band is formed that is resistant to I-*Sce*I digest *in vitro*. **(B)** Genomic blots. The colored line above the blots indicates the genotype, WT (W11579) cells in black and *mre11Δ* (W11580) in green. The blots on the left show the appearance of the 7 kb I-*Sce*I-cut band (cut) after *in vivo* DSB induction at the I-*Sce*I cut site. The blots on the right show the same DNA samples from the left after I- *Sce*I digestion *in vitro*. The 8 kb I-*Sce*I-resistant repair product results from gene conversion (GC). **(C)** Quantification of cutting and repair products was performed on the blots shown in B using ImageJ and are depicted as the percent of total signal for each lane. The percent I-*Sce*I cutting and gene conversion were calculated using the equations above the graph for WT and *mre11Δ* cells. Error bars represent the standard deviation from 3 biological replicates. Note that the 30’ and 60’ time points shown in the graph are not shown in the genomic blots. In addition, the 18 kb *lacO* fragment is not shown.

### Overexpression of Dna2 suppresses the delay in HR events in *mre11Δ* cells

Thus far, we have shown that a temporal relationship exists between DNA end processing, recruitment of HR proteins, chromosome mobility, homolog pairing, and gene conversion. Since all of these events are delayed in *mre11Δ* cells, we searched for a suppressor of the *mre11Δ* resection delay and asked whether it could concomitantly rescue downstream events. Since overexpression of Exo1 suppresses *mre11Δ* γ-ray sensitivity in haploid cells (Lewis, Karthikeyan, Westmoreland, & Resnick, 2002; Mimitou & Symington, 2010; Moreau et al., 2001; Tsubouchi & Ogawa, 2000), we first examined whether Exo1 overexpression could rescue DNA end resection and the timing of gene conversion in *mre11Δ* cells. Although overexpression of Exo1 rescues γ-ray sensitivity in both haploid and diploid *mre11Δ* cells (Fig. S4), surprisingly, it does not rescue the *mre11Δ* delay in resection, Rad51 protein recruitment, increased mobility or gene conversion (Fig. 5, brown lines). In contrast, overexpression of Dna2, another nuclease that plays an important role in DNA end resection (Niu et al., 2010; Symington, 2016; Zhu et al., 2008) does not rescue γ-ray sensitivity of either haploid or diploid *mre11Δ* cells (Fig. S4). However, Dna2 overexpression increases the percentage of ssDNA 0.3 kb from the DSB (Fig. 5A, turquoise vs. green), shifts the timing of Rad51 recruitment to a DSB (Fig. 5B, turquoise vs. green). rescues chromosome mobility 3 hours after induction to near WT levels (Fig. 5C, turquoise vs green and Fig. 1C) and rescues the timing of gene conversion (Fig. 5D, turquoise vs green). Taken together, these data demonstrate that, upon Dna2 overexpression, the strong temporal linkage between DNA end resection, chromosome mobility, and gene conversion move in concert and depend on the initial timing of DNA end processing. These observations not only highlight the importance of ssDNA formation as a vital precursor to increased chromosome mobility, but also support the hypothesis that mobility and HR are mechanistically linked.

**Figure 5.**
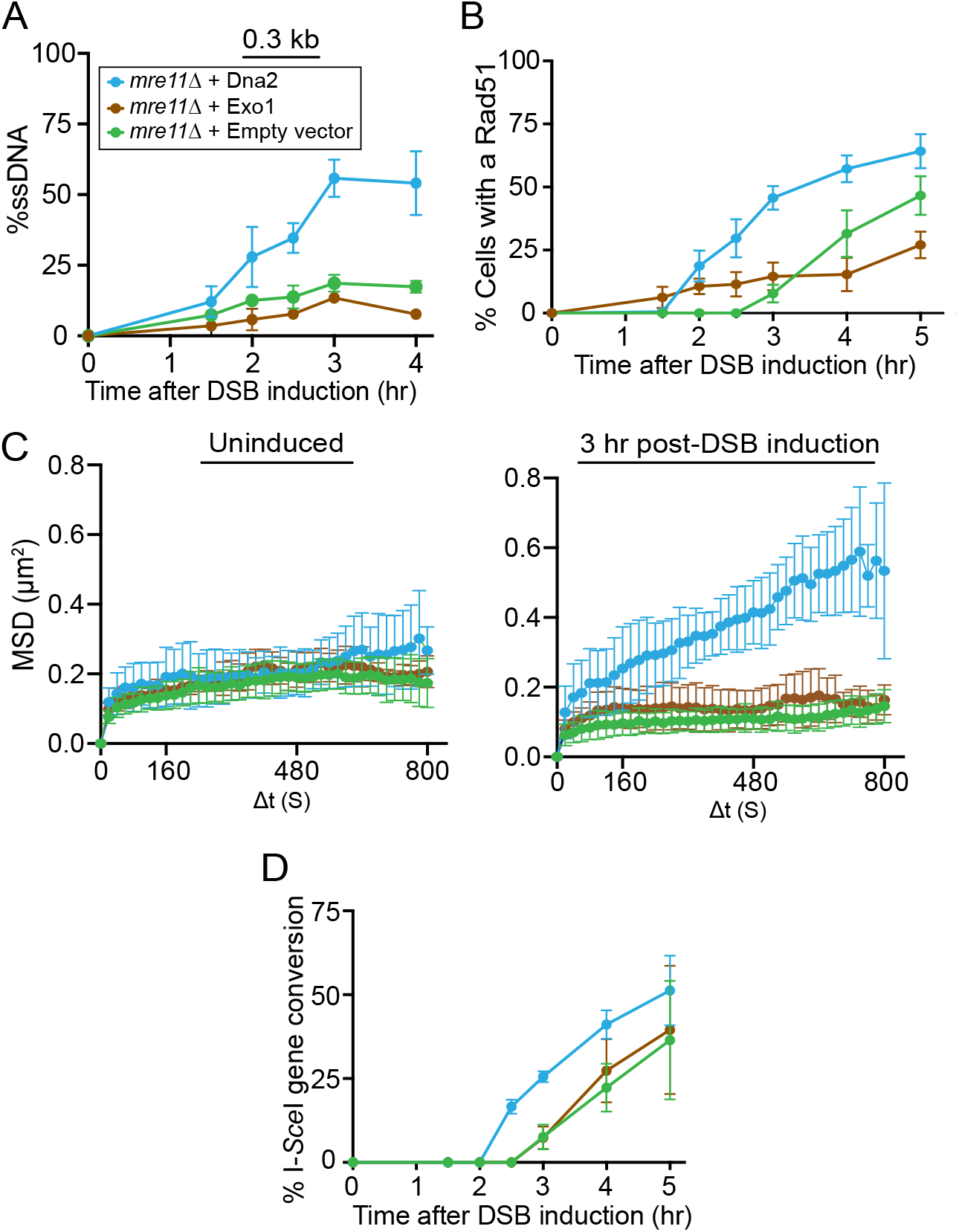
Overexpression of Dna2 suppresses the delay in HR events in *mre11Δ* cells. **(A)** Percentage of ssDNA present at the 0.3 kb *Dra*I sites was measured by the qPCR assay described in Figure 2 and plotted vs. time after DSB induction. Shown are *mre11Δ* + empty vector (U3377), *mre11Δ* + Dna2 (U3376), and *mre11Δ* + Exo1 (U3375) cells in green, cyan, or brown respectively. **(B)** Rad51 recruitment to the I-*Sce*I cut site after DSB induction in *mre11Δ* + empty vector (U3426), *mre11Δ* + Dna2 (U3427), and *mre11Δ* + Exo1 (U3428) cells. At each time point, at least 100 cells were analyzed for each strain. **(C)** MSD plot for the *tetO* array in *mre11Δ*+ empty vector (U3429), *mre11Δ* + Dna2 (U3430), and *mre11Δ* + Exo1 (U3431) cells. On the left cells were untreated (n = 10, 12, 11) or on the right induced for 4 hours of I-*Sce*I cutting (n = 10, 9, 8) respectively. Error bars of MSD plots represent the 95% CI. **(D)** Gene conversion as measured by genomic blot assay. Cells were transformed with a galactose-inducible overexpression plasmid and assessed for gene conversion after I-*Sce*I DSB induction. Percent gene conversion vs. time is plotted for the same strains shown in (A). Error bars represent the standard deviation from 3 biological replicate blots of 3 individual transformants.

## Discussion

In this study, we demonstrate that changes in the timing of chromosome mobility result in a corresponding change in essential downstream HR events. Mutations in genes involved in the DNA damage response, chromatin remodeling, and HR repair eliminate increased chromosome mobility (Dion et al., 2012; Mine-Hattab & Rothstein, 2012; Seeber et al., 2013). However, it is not clear whether or how much of the reduction in DNA repair can be attributed solely to the loss of chromosome mobility versus the loss of recombination. To explore the relationship between chromosome mobility and HR, we examined key processes in the HR pathway at multiple time points after DSB induction in WT cells and a mutant that alters the timing of chromosome mobility. We find that DNA end processing, recombination protein loading, chromosome mobility, physical pairing of homologs and gene conversion are all temporally linked. These events all move together whether mobility is slowed in the mutant cell or accelerated after overexpression of a nuclease (Fig. 6).

**Figure 6.**
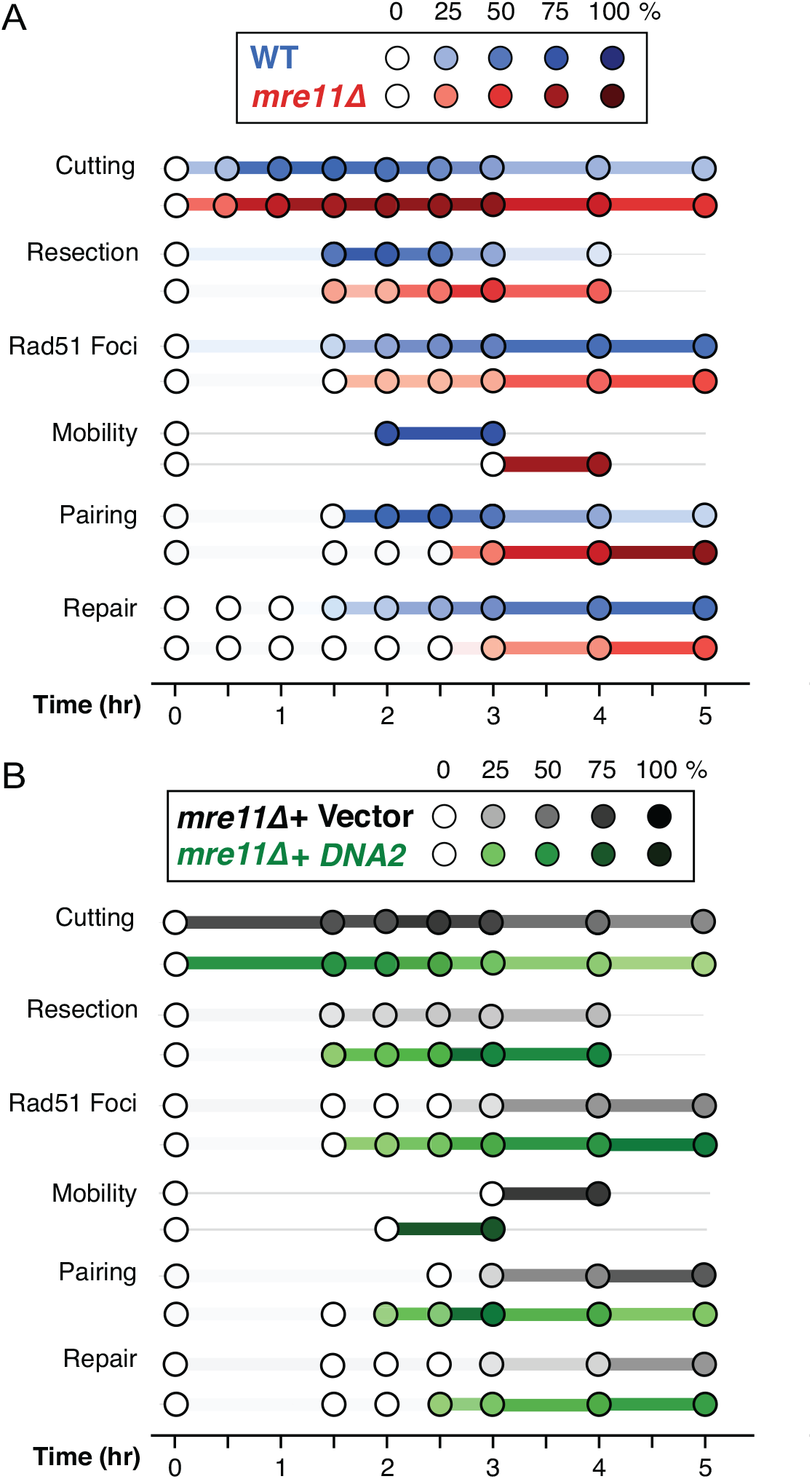
Chromosome mobility and HR are linked. **(A)** Summary plots with the concordance of HR events examined in this study on the Y-axis and time after DSB induction on the X-axis (values listed in Table S3). WT cells are shown in blue and *mre11Δ* cells in red. The color scale from light to dark represents the percent of each event measured. The time after DSB induction that results in the maximum measurement in each time series is set to 100%. Specifically, for homolog pairing, the 5% background that occurs in the absence of recombination (Fig. 3, *rad51Δ*) is subtracted from each time point. **(B)** Summary plots showing the various HR events for *mre11Δ* + empty vector in black and *mre11Δ* + Dna2 in green, as described in (A) (values listed in Table S3).

Although we show that mobility is linked to HR in yeast, some groups have argued that increased chromosome mobility is not required for HR (C. S. Lee et al., 2016; Strecker et al., 2016). In those studies, mobility was only examined at a single time point, and repair outcomes were assessed via colony growth 3-4 days post DSB induction, leaving a large time frame in which increased mobility could have occurred. In our study, the events of DSB repair are followed over several hours after break induction, allowing a much closer look into the window where changes in chromosome mobility likely play an important role.

Our study also shows that the formation of ssDNA is an important regulator of increased chromosome mobility, similar to what was observed in mammalian cells, where inhibiting the nuclease activity of MRN reduces end resection and substantially decreases the movements of DSBs (Schrank et al., 2018). However, a recent report argues that mobility is not correlated with the efficiency of DNA end resection in yeast (Cheblal et al., 2020). As we show here with *mre11Δ*, as well as in our previous study with *sae2Δ* (Mine-Hattab & Rothstein, 2012), it is critical to know the time of DSB induction relative to the time of the mobility assay. For example, increased chromosome mobility is delayed in *mre11Δ* cells and would be missed if measured too early. Similarly, the reverse could also be true: if DNA end resection occurs earlier, then increased chromosome mobility would also occur early and would be missed if assessed too late. Thus, our observation that delayed HR events in an *mre11Δ* mutant can be suppressed by Dna2 overexpression (Fig. 6A, B) and our studies of resection (Fig. 2 C, D and Fig. 5A) further support the link between ssDNA formation and chromosome mobility.

It is surprising that Exo1 overexpression, which suppresses *mre11Δ* γ-ray sensitivity (Lewis et al., 2002; Moreau et al., 2001; Tsubouchi & Ogawa, 2000) (Fig. S4), is unable to rescue the delayed timing of HR events in *mre11Δ* cells induced by a single DSB (Fig. S6). In contrast, overexpression of Dna2, which is also involved in end resection, does not suppress γ-ray sensitivity (Fig. S4), but rescues the *mre11Δ* delay in mobility and the subsequent downstream HR events after an I-*Sce*I-induced break (Fig. 6B). Perhaps this difference is due to the way these two nucleases interact with the Ku complex (yKu70/80) (Fig. S7, left). When Ku is bound to the ends of a DSB *in vitro*, it specifically blocks Exo1 access, but is less effective at blocking the recruitment of Dna2 (Shim et al., 2010). Consistent with these *in vitro* findings, resection is reduced in *mre11Δ* cells, where the Ku complex no longer directly competes with the MRX complex for free DNA ends, effectively blocking Exo1 entry (Mimitou & Symington, 2010; Shim et al., 2010; Tomita et al., 2003). Another critical difference between these two nucleases relates to their interactions with nucleosomes (Fig. S7, right). Resection by Exo1 is blocked by nucleosomes, whereas resection by the Sgs1-Dna2 complex actively displaces them during its movement along DNA (Adkins, Niu, Sung, & Peterson, 2013; Xue et al., 2019). In WT cells, histone removal and chromatin remodeling occur at the site of a DSB as a downstream consequence of both end processing and DNA damage checkpoint activation initiated by the MRX complex (Hauer et al., 2017; Seeber et al., 2013). In *mre11Δ* cells, we suspect that there is insufficient checkpoint activation after induction of a single DSB to remove histones and allow Exo1 to efficiently resect DNA. On the other hand, Dna2 does not require chromatin remodeling for efficient resection at a break site (Adkins et al., 2013). Ultimately, the differences between these two nucleases requires further investigation.

Unanswered questions also remain surrounding the mechanism causing increased chromosome mobility after DSB formation. Given that the recombination protein Rad51 is essential for chromosome mobility (Dion et al., 2012; Mine-Hattab & Rothstein, 2012; Smith & Rothstein, 2017), it has been proposed that its recruitment to resected DNA ends results in a stiff filament that allows the navigation of the broken DNA end through the densely packed chromatin in the nucleus (Mine-Hattab, Recamier, Izeddin, Rothstein, & Darzacq, 2017). Other models propose that in response to DNA damage, checkpoint activation and chromatin remodeling cause changes in chromatin stiffness, which promote increased chromosome mobility (Hauer et al., 2017; Herbert et al., 2017; Seeber & Gasser, 2017). Additionally, changes in chromosomal conformation due to H2A phosphorylation have been proposed to be sufficient for increased chromatin mobility, even in the absence of DNA damage (Garcia Fernandez et al., 2021). Our finding that DNA end resection is a vital precursor to increased chromosome mobility is compatible with the above models, since RPA-bound ssDNA enhances checkpoint activation (Zou & Elledge, 2003), which is required for H2A phosphorylation (Rossetto, Avvakumov, & Cote, 2012; Shroff et al., 2004) and aids in the recruitment of chromatin remodeling proteins (Eapen, Sugawara, Tsabar, Wu, & Haber, 2012; van Attikum, Fritsch, & Gasser, 2007). RPA-bound ssDNA is also required for Rad51 presynaptic filament formation (Sung, 1994; Sung & Robberson, 1995). Thus, aspects of these different models are not mutually exclusive and, in fact, checkpoint activation, H2A phosphorylation, chromatin remodeling and Rad51 filament formation likely work in concert to produce the changes in chromosomal motion necessary for genetic recombination between homologs (Fig. 7). In summary, the tight temporal relationship we observe here between DNA end resection, homolog pairing, and gene conversion along with our inability to separate mobility from DNA repair, strongly supports the idea that increased chromosome mobility is mechanistically linked to HR.

**Figure 7.**
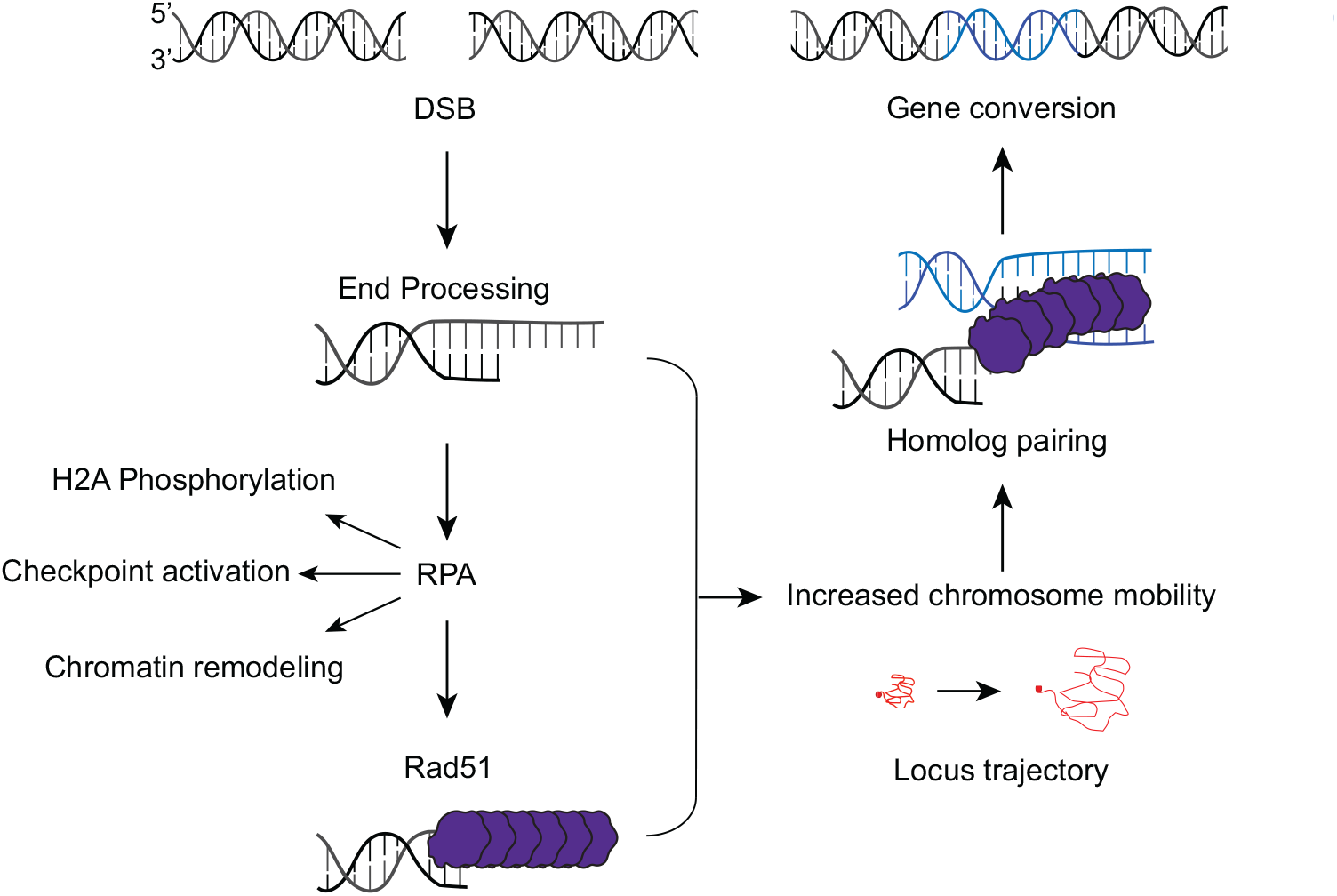
Pathway to increased chromosome mobility and HR. Diagram showing the different steps that occur after DSB formation that contribute to increased chromosome mobility. After a DSB is formed, the DNA ends are processed to create 3’ ssDNA tails. We show here that the timing of end processing is a vital precursor for the initiation of chromosome mobility. Additionally, the ssDNA formed binds RPA, which enhances checkpoint activation, is required for H2A phosphorylation, and aids in the recruitment of chromatin remodeling proteins. RPA-bound ssDNA is also required for Rad51 presynaptic filament formation, since Rad51 loading occurs via the physical exchange with RPA through the mediator Rad52 (not shown). All of these events contribute to increased chromosome mobility, measured by tracking the locus trajectory at the site of a DSB (denoted in red). The increase in chromosome mobility leads to homolog pairing and ultimately repair of the break via gene conversion.

## Methods

### Strains and plasmids

All strains are *RAD5+* derivatives of W303 (Thomas & Rothstein, 1989; Zhao, Muller, & Rothstein, 1998) (see Table S1). Plasmid pWJ2216 was constructed to express genes bidirectionally from either the *pGAL1* or *pGAL10* promoters. The *TEF* terminator sequence from plasmid pAG36 (Goldstein & McCusker, 1999) was PCR-amplified and cloned into the *Nru*I cut site adjacent to the transcription start site for the *GAL10* promoter of plasmid pWJ1781 (Reid et al., 2016). Terminator sequence cloning reestablished the *Nru*I cut site proximal to the *GAL10* promoter. *DNA2* was amplified with forward (gacgaaagggcctcgtgtcgATGCCCGGAACGCCACAGAAGAAC) and reverse (gaatctttttattgtcagttcgTCAACTTTCATACTCTTGTAGAATTTCC) primers that include 5’ tails (lower case text) to allow cloning by recombination with the *Nru*1-cut plasmid directly in yeast to form plasmid pWJ2563. *EXO1* was amplified in a similar way with forward (gacgaaagggcctcgtgtcgGCTCGAAAAAACTGAAAGGCGT) and reverse (gaatctttttattgtcagttcgCCTCCGATATGAAACGTGCAGTA) primers and cloned to produce plasmid pWJ2620.

### Media and galactose induction

Rich medium (yeast extract–peptone–dextrose, YPD) was used as described previously (Sherman et al, 1986). For synthetic complete (SC) medium lacking the appropriate amino acids, a custom synthetic complete amino acid mixture with 2× L-leucine was used as described in (Sherman, Fink, & Hicks, 1987). For I-*Sce*I DSB induction, cells were grown overnight at 23°C in 7 ml of SC lactate liquid medium (Synthetic complete amino acid mixture; 3% glycerol; 2% lactic acid; 0.05% glucose; 150 mg/L adenine; adjusted to pH 5.7 with sodium hydroxide) in sterile 15 ml culture tubes (Joseph, Lee, Bryant, & Rothstein, 2021). The following morning, the cultures were diluted in 7 ml of fresh SC glycerol lactate medium to an optical density at a 600 nm wavelength (OD_600_) of approximately 0.3. Then the cultures were allowed to grow for 2 to 3 hours or until they reached approximately 0.5 OD_600_. To start the DSB induction, galactose was added to the cultures at a final concentration of 2%. Next, 1 ml of the growing cultures were removed and placed into a separate 1.5-ml centrifuge tube at each of the desired time points of DSB induction. Cells were then prepared for microscopy as described below.

### Microscopy

For each of the collected time points after DSB induction, the cells were used immediately to prepare slides for image acquisition. Cells were pelleted by centrifugation and then washed in SC glucose medium to repress I-*Sce*I expression. 3 µl of the resuspended cells were then mixed with 3 µl of 1.4% melted agarose and placed on the center of a glass microscope slide. A glass coverslip was placed on top of the cell suspension and slide edges were sealed with melted wax. Image acquisition was performed using an epifluorescence microscope, the Leica DM5500B upright microscope (Leica Microsystems), illuminated with a 100W mercury arc lamp or a Prizmatix© LED light and captured with a Hamamatsu Orca AG cooled digital CCD (charged- coupled device). We used a 100x objective and high-efficiency filter cubes for fluorophore imaging (Chroma 41028, Chroma 31044v2, and Chroma 41002C, for YFP, CFP, and RFP, respectively). The fluorophores used in our diploid strains are the red- and blue-shifted variants of GFP, YFP(10C) and CFP(W7), respectively (Heim & Tsien, 1996; Ormo et al., 1996), as well as the monomeric version of DsRed (mRFP1) (Campbell et al., 2002). The peak emission wavelength used for the detection of YFP, CFP and RFP are 535 nm, 480 nm and 620 nm, respectively. To ensure that cells were in early S-phase, only small-budded cells were selected for imaging.

### Image analysis

Image analysis was performed with Volocity software (Perkin-Elmer). Time-lapse images were identically deconvolved using the iteration deconvolution tool in the Volocity software for contrast enhancement. The find objects tool of the Volocity software was used to track the three-dimensional (3D) positions of the tagged loci and identify the intensity-weighted mass center for each focus. A fluorescence intensity of 10 standard deviation above background was used as a cut off for the *lacO*/LacI–YFP and *tetO*/TetR–RFP arrays.

### Chromosome mobility

To measure chromosome mobility, a mean square displacement (MSD) analysis (Gasser, 2002; Marshall et al., 1997; Mine-Hattab & Rothstein, 2012) was used to track the change in displacement of a fluorescently tagged locus near the site of an inducible DSB. To calculate the MSD values, time-lapse movies were taken of the cells and the 3D positions of the *tetO* array (RFP focus) and the spindle pole body (Spc110–YFP focus) were recorded every 20 seconds. To correct for the drift due to the motion of the nucleus inside the cell, the position of the tetO locus was subtracted from the position of the Spc110 focus using the equation (x_tetO_–x_Spc110_, y_tetO_ – y_Spc110_, z_tetO_ –z_Spc110_). We utilized an R script to measure the MSD of the translocations of the tetO locus in each movie using the equation ti = < (xi+1 −xi)2 +(yi+1 −yi)2 +(zi+1 −zi)2 >. These values were the plotted over the change in time (Δt). The radius of confinement (R_c_) was then calculated by averaging the individual MSD plots for each cell in an experiment into a mean value, fitting the resultant curve, and taking ¾ of value the plateau of that curve. Due to photo bleaching, we could not image longer than 20-minutes before the fluorescence signal could no longer be distinguished from background. A previous study from our lab shows that MSD plots plateau between 1100 and 1200 seconds after induction of I-*Sce*I cutting (Mine-Hattab & Rothstein, 2012). In the study presented here, the maximal time interval is limited to 900 seconds due to photobleaching. Therefore displacement, at 900 seconds is used as a minimal estimate of the plateau.

### Homolog pairing

Homolog pairing was performed as previously described in Joseph, et al. (2021). Briefly, cells were grown overnight in 7 ml of SC lactate liquid medium, re-diluted the next morning to an OD_600_ of 0.3 and allowed to grow at 23°C until they reached an OD_600_ of approximately 0.5. Next, the culture was divided into separate 1 ml tubes and galactose was added to a final concentration of 2% to induce a DSB at the I-*Sce*I cut-site. After 0, 1.5, 2, 2.5, 3, 4, and 5 hours of DSB induction each culture was prepared for cell imaging as described in the microscopy section above. Volocity software (Perkin-Elmer) was used to image and identify small-budded S-phase cells and their fluorescently labeled homologs. For each cell acquired, the X,Y, and Z positions of the homologs was recorded and the distance between them measured. The percent of homolog pairing at each time point of DSB induction was calculated using the pairing threshold that was empirically determined as described in Fig S2.

### Genomic blots

To quantify the amount of I-*Sce*I cutting and I-*Sce*I gene conversion after DSB induction, cells were grown overnight in 25 ml of SC glycerol lactate liquid medium and re-diluted the next morning in 50 ml of the same medium for 3 hours until they reached an OD_600_ of approximately 0.5. Next, galactose was added to a final concentration of 2% to induce a DSB at the I-*Sce*I cut-site. After 0, 30, 90, 120, 150, 180, 240, and 300 min of DSB induction in galactose, 7 ml of the growing culture was sampled, and DNA was extracted. To quantify the amount of I-*Sce*I cutting, 5 µg of DNA from each time point was digested with *Sgr*AI and *Xma*I to cut chromosome V adjacent to the I*-Sce*I cut-site and in the *tetO* array, respectively. Uncut chromosomes containing the *tetO* array produce an 8 kb fragment, whereas I-*Sce*I-cut chromosomes produce a 7 kb and a 1 kb fragment (Fig. 5A). To quantify the amount of I-*Sce*I gene conversion, 5 µg of DNA from the same extract was digested *in vitro* with *Xma*I, *Sgr*AI, and purified I-*Sce*I. Only *tetO* array containing chromosomes that have repaired the I-*Sce*I cut-site via gene conversion using the homologous site on the *lacO* array-containing chromosome V are I-*Sce*I resistant and thus produce an 8 kb fragment. DNA was then loaded on a TBE + 0.4% agarose gel, run overnight at 25 volts, transferred to a membrane, crosslinked, and labeled with a 1 kb radioactive probe in the *GEA2* gene (primers shown in Table S2). The probe is located between the I-*Sce*I site and the *tetO* array (Fig. 5A).

### qPCR-based resection assay

To measure resection by qPCR, we use an approach first described by Zierhut and Diffley (2008) as modified by Gnügge et al. (2018). In this approach, we chose genomic DNA sequences containing *Dra*I restriction sites located at various distances from the I-*Sce*I-induced DSB (Fig. 2A). PCR primers were designed to amply those sites. The *Dra*I endonuclease can only digest DNA that is double stranded, thus *Dra*I digestion prior to PCR prevents amplification of those sites. However, if resection has started at the induced I-*Sce*I DSB and has passed the *Dra*I restriction site, that site becomes single stranded and resistant to *Dra*I digestion, thus allowing PCR amplification (Fig. 2A). As described previously, for each time point, the percent ssDNA is calculated using the following equation: ssDNA% = 2/(2^(ΔCt)+1)*100 (Gnügge et al., 2018). To correct for differences in template concentrations, an amplicon in the *ADH1* gene is used as a control site to normalize the Ct values obtained for the *DraI* sites. The qPCR assay is performed in a 384-well format using the Bio-Rad SsoAdvanced™ Universal SYBR^®^ Green Mix on a Bio-Rad CFX384™ Real-Time System.

### Plate assay for radiation sensitivity

Haploid or diploid cells were grown in liquid SC-Leu medium to mid-log phase. The cultures were fivefold serially diluted and spotted onto solid YPD or YPGal plates. The plates were placed in a Gammacell-220 irradiator containing ^60^Co as the irradiation source and exposed to 40, 100, or 200 Gy of γ-irradiation. After irradiation, the plates were incubated for 2 days at 30°C before being imaged. WT or *mre11Δ* cells were transformed with a galactose-inducible overexpression plasmid [empty vector (EV), Exo1 or Dna2].

## Acknowledgments

We thank the entire Rothstein laboratory including, Michael Smith, Olga Marte, Gaël Fortin, and Keerthana Chetlapalli for experimental feedback and suggestions. We also thank Judith Miné-Hattab and Michael Lisby for their work in the initial construction of the yeast strains used in this study. We thank Lorraine Symington, Roberto Donnianni, Robert Gnuegge, and Michael Kimble for their feedback and help with genomic blotting and qPCR. Lastly, we also thank Eric Greene, Alberto Ciccia, and especially Luke Berchowitz for careful reading of the manuscript. This work was supported by National Institutes of Health grants T32 GM007088 (to F.J.J.), T32 CA009503 (to F.J.J. and E.E.B.), R35 GM118180-S1 (to F.J.J.), T32 GM008798 (to E.E.B.), TL1 TR001875 (to E.E.B.), and R35 GM118180 (to R.R.).

---

**Figure S1.**
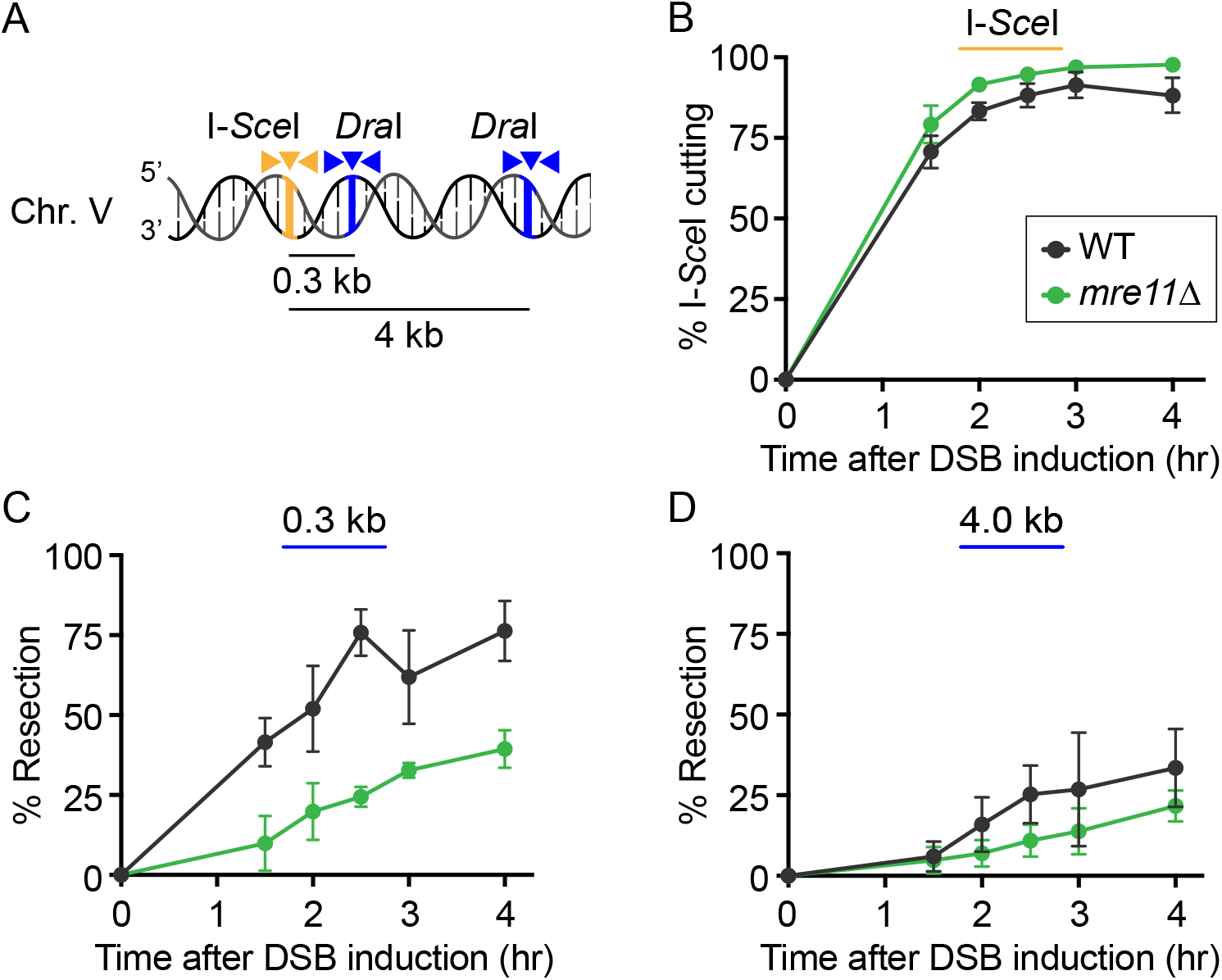
DNA end-resection is delayed in haploid *mre11Δ* cells. **(A)** Schematic of the haploid strain used to measure ssDNA via qPCR. *Dra*I sites (blue lines) are located either 0.3 kb or 4.0 kb away from the I-*Sce*I cut site (yellow line). PCR primers that flank each *Dra*I site (blue arrowheads) are used for amplification before and after *in vitro* digestion with *Dra*I. The percent of *Dra*I-resistant DNA reveals the amount of DNA resection at that site. Amplification of DNA from primers that flank the I-*Sce*I site (yellow arrowheads) is used to measure I-*Sce*I cutting. **(B)** Percentage of I-*Sce*I cutting plotted vs. time after DSB induction. **(C-D)** Percentage of ssDNA present at the 0.3 kb and 4.0 kb *Dra*I sites vs. time after DSB induction. In all graphs, WT (W9561- 17A) is shown in black and *mre11Δ* (W11027-14B) in green. Error bars represent the standard deviation from 3 biological replicates.

**Figure S2.**
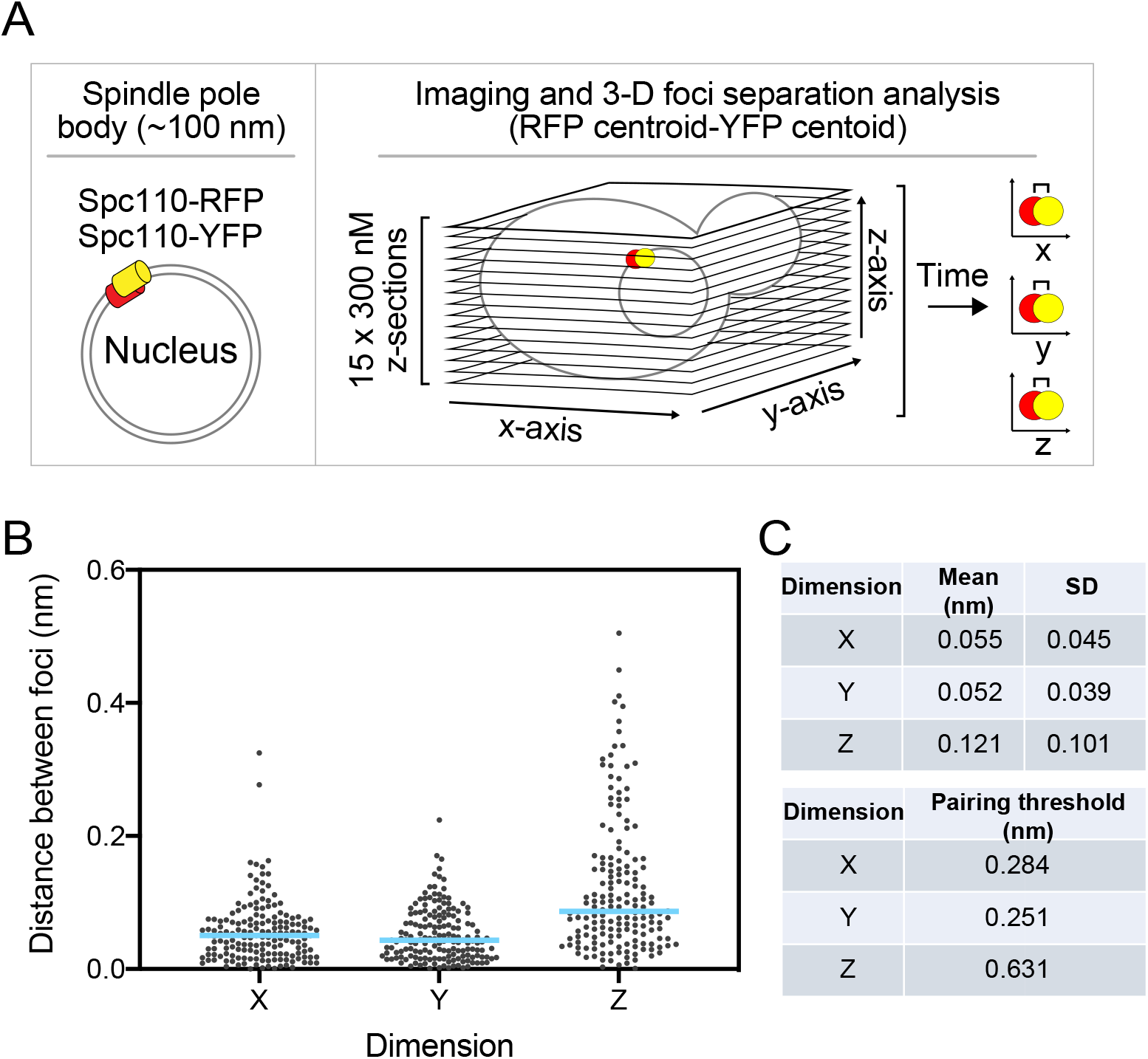
Determining the threshold for homolog pairing. To properly define a threshold for homolog pairing and account for errors in microscope precision, we performed a calibration experiment. **(A)** Schematic of the strain used to determine the threshold for homolog pairing in diploid yeast cells. The two homologous copies of the spindle pole body component, Spc110, are tagged with YFP and RFP, respectively. The resulting cells display a single focus on the nuclear periphery labeled with the same two fluorescent tags used to detect the two homologs (see Fig. 3). Ideally the distance between the centroid position of the red and yellow Spc110 foci should be zero if there is no error in measurement or image acquisition. **(B)** The measured distance between the centroid positions of the two foci in a given cell is plotted for the X, Y, and Z dimension. Each black dot represents one cell (n = 266). The blue line represents the mean. **(C)** The mean distance between the two foci across all cells measured and the variance of the mean in each dimension is shown in the top panel. To maximize recovery of true positives while minimizing false positives, a range of 5 standard deviations within the mean distance observed in each dimension was chosen (∼99% positive recovery rate). The threshold used for homolog pairing is shown in the bottom panel. Two foci are considered paired when the distance between them is at or below the threshold for each dimension.

**Figure S3.**
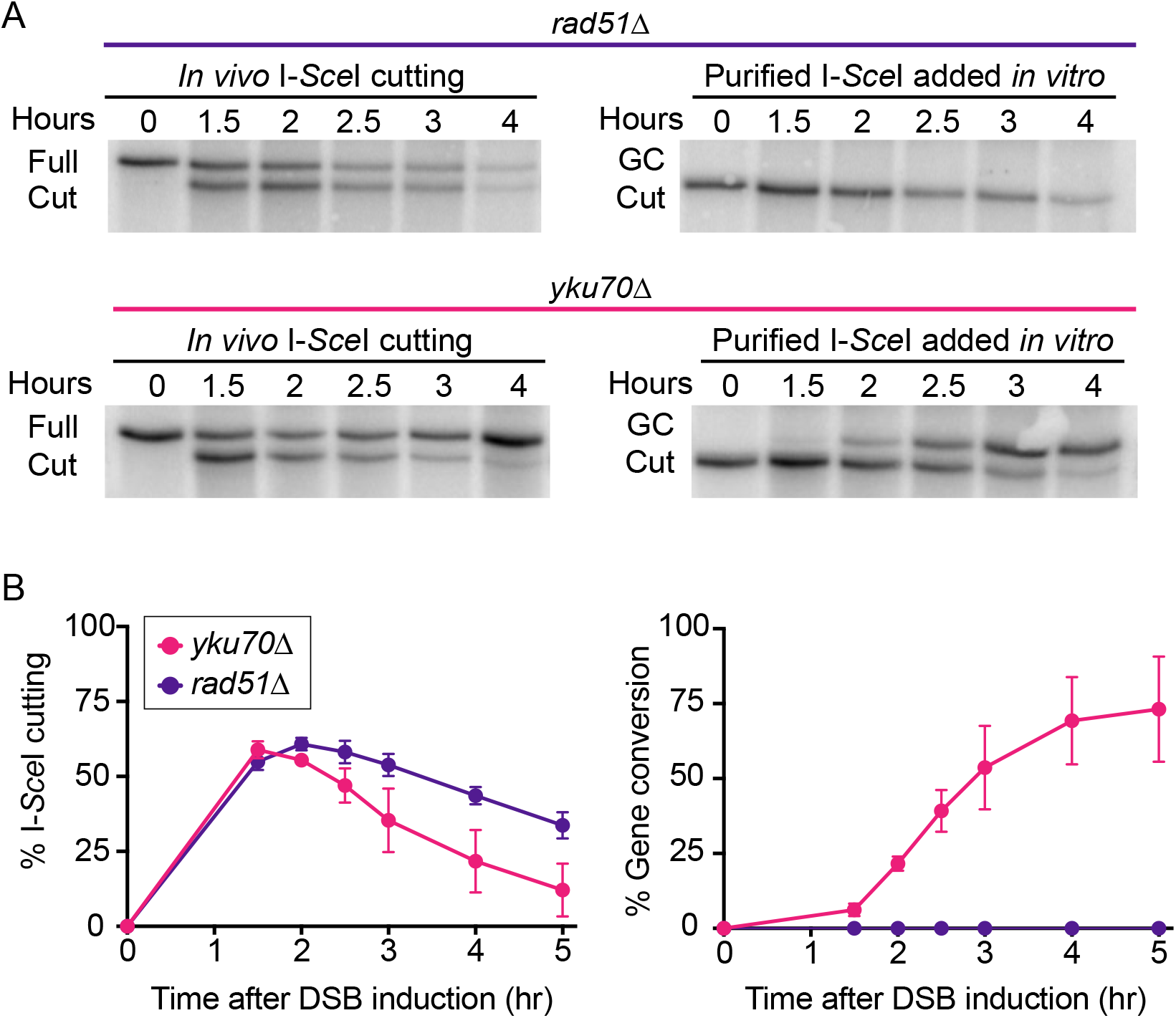
Gene conversion is dependent on the Rad51 recombinase and independent of NHEJ. **(A)** Genomic blots. The colored line above the blots indicates the genotype, *rad51Δ* (W11585) in purple and *yku70Δ* (W11586) in pink. The blots on the left show the appearance of the 7 kb I-*Sce*I-cut band (cut) after *in vivo* DSB induction at the I-*Sce*I cut site. The blots on the right show the same DNA samples from left but after *in vitro* digestion with purified I-*Sce*I enzyme. The 8 kb I-*Sce*I-resistant repair product results from gene conversion (GC). **(B)** Quantification of cutting and repair products was performed using ImageJ and are shown as the percent of total signal for each lane. The percent I-*Sce*I cutting and gene conversion were calculated for *rad51Δ* and *yku70Δ* as described in Figure 4. Error bars represent the standard deviation from 3 biological replicates. Both *in vivo* I-*Sce*I cutting and the appearance of the 8 kb I-*Sce*I-resistant band were unaffected by Yku70 deletion, demonstrating that NHEJ plays no significant role in producing the I-*Sce*I resistant product detected in our assay. Furthermore, the appearance of the 8 kb I-*Sce*I-resistant band is dependent on the Rad51 recombinase, supports the notion that this band is the product of gene conversion.

**Figure S4.**
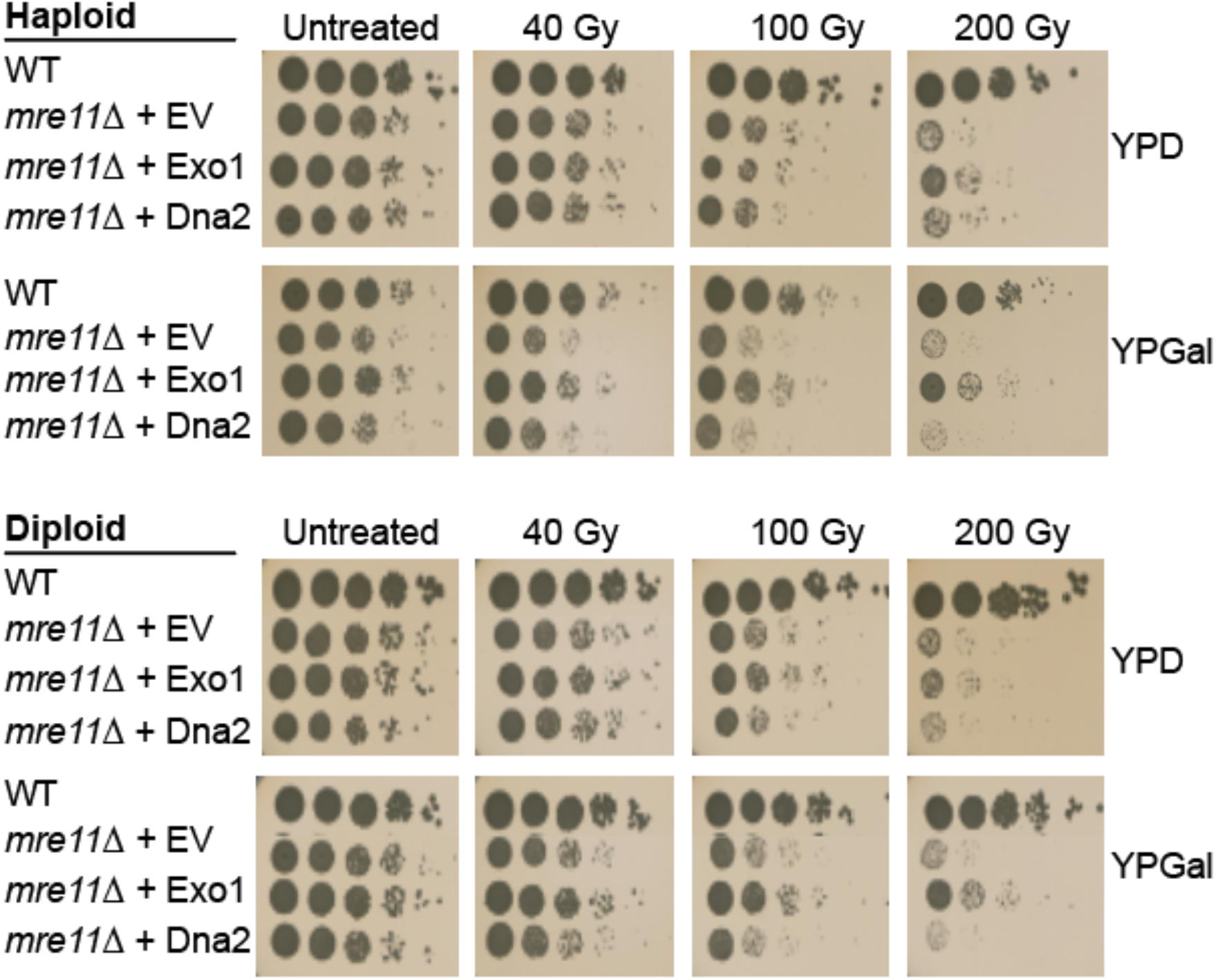
Overexpression of Exo1 suppresses *mre11Δ* γ-ray sensitivity in both haploid and diploid cells. Spot assay to assess DNA damage sensitivity. Fivefold serial dilutions of haploid (top) and diploid (bottom) cells grown for 2 days after exposure to 40, 100, or 200 Gy of γ-irradiation. Cells are either WT or *mre11Δ,* transformed with a yeast centromere galactose-inducible overexpression plasmid (empty vector (EV), Exo1 or Dna2). Growth of the strains on YPD and YPGal plates before and after γ-irradiation is shown.

**Figure S5.**
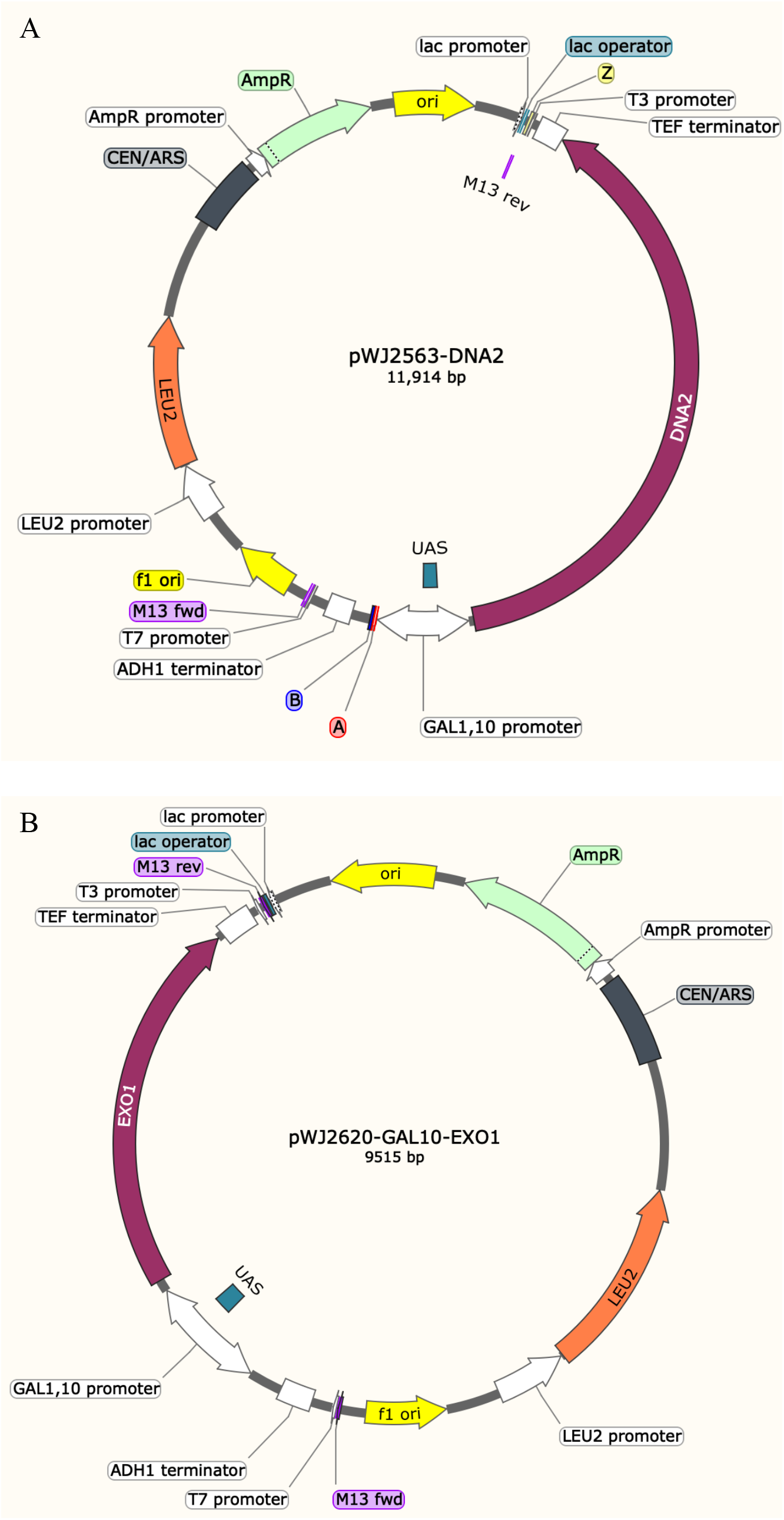
Map of plasmids pWJ2563 and pWJ2620, used for overexpressing Dna2 and Exo1. **(A)** The *DNA2* gene coding sequence was PCR- amplified from yeast genomic DNA and cloned by *in vivo* recombination with the C and D adaptamer sequences into a *Nru*l digested plasmid to produce the plasmid pWJ2563 as described in (Reid et al., 2016). *In vivo* recombination places *DNA2* downstream of a *GAL110* promoter for galactose inducible expression. *LEU2* is used as the selectable maker for the vector under the control of its native promoter. **(B)** The *EXO1* gene was similarly cloned into the same vector as in (A) to produce the plasmid pWJ2620. The diagrams representing the plasmids were both created using Snap gene® software and are not to scale. Plasmids were verified using DNA sequencing and are available upon request (see Table S2).

**Figure S6.**
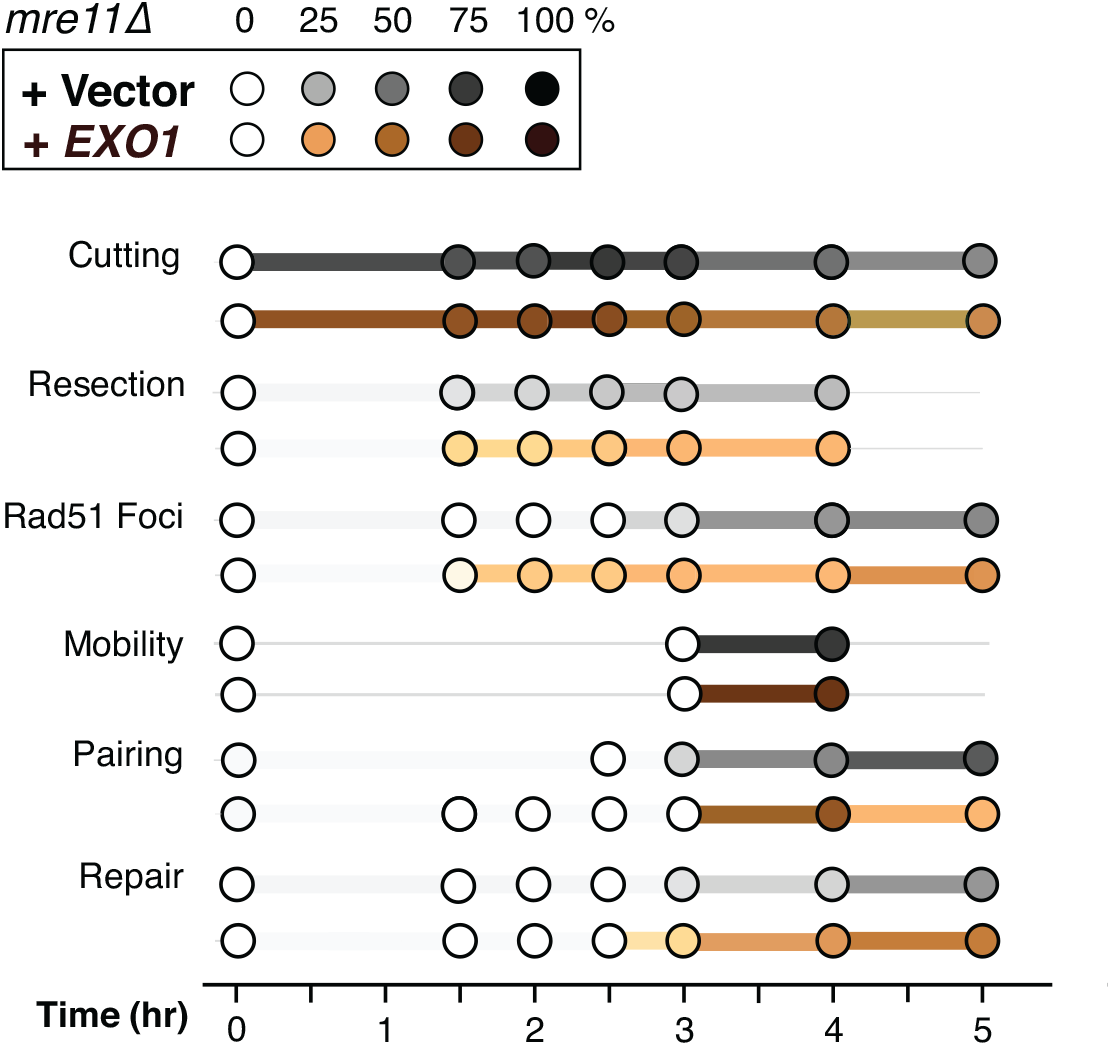
Exo1 overexpression does not suppress the delay in HR events in *mre11Δ* cells. Summary plots showing the various HR events for *mre11Δ* + empty vector in black and *mre11Δ* + Exo1 in brown (values listed in Table S3). The color scale from light to dark represents the percent of each event measured. The time after DSB induction that results in the maximum measurement in each time series is set to 100%. Specifically, for homolog pairing, the 5% background that occurs in the absence of recombination is subtracted from each time point.

**Figure S7.**
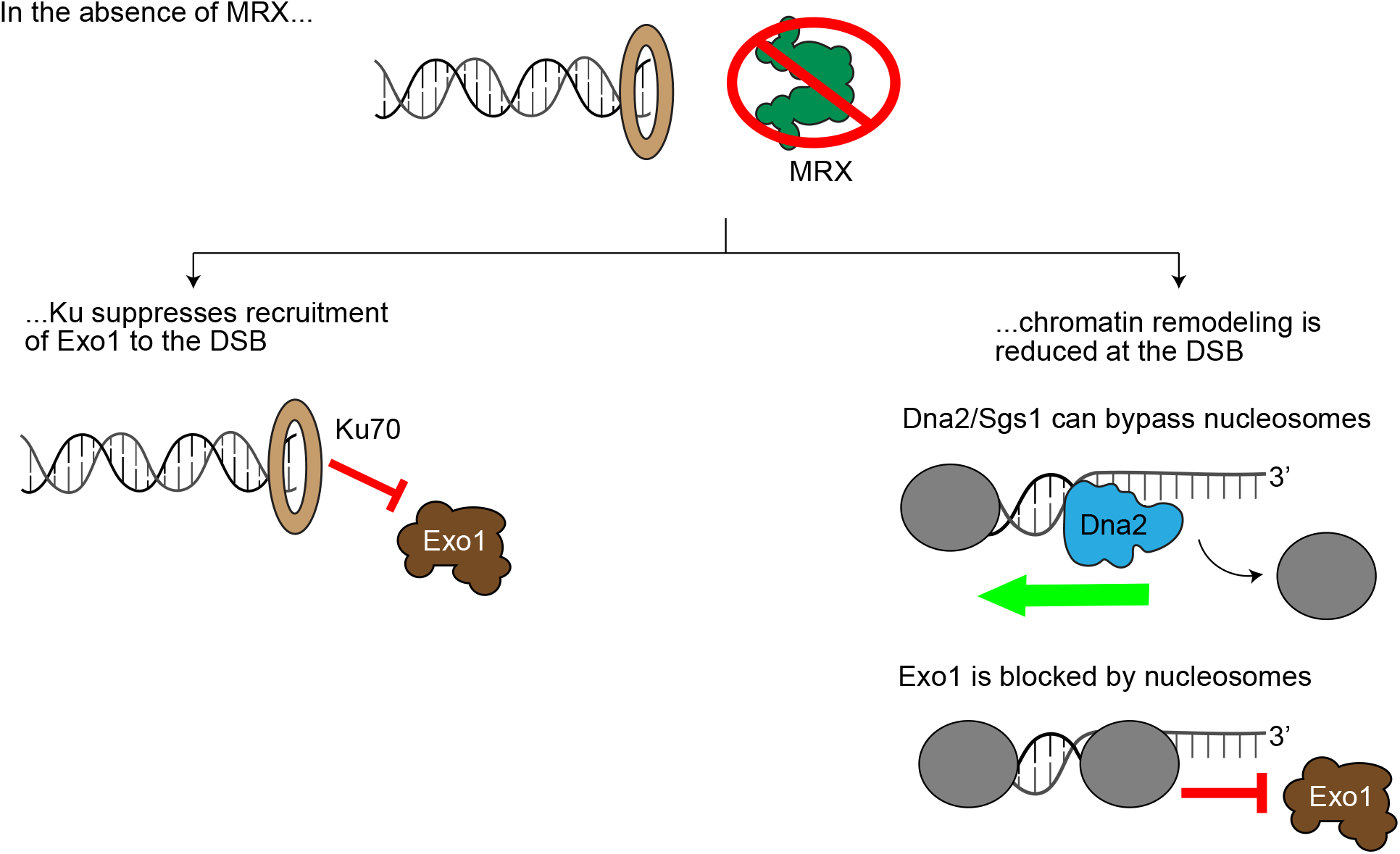
Key differences between Dna2 and Exo1. When a DSB is formed in a WT cell, both MRX and Ku bind the broken ends. MRX acts to remove Ku and help recruit the nucleases Dna2 and Exo1 to the break site. However, in the absence of MRX, Ku is more abundant and specifically suppresses Exo1 (left). Additionally, genomic DNA in the cell is not naked but bound by histones and packaged into nucleosomes. Dna2 and the helicase Sgs1 (not shown) can act together to unwind DNA and bypass nucleosomes, whereas Exo1 cannot and is impeded by their presence (Wright et al.).

**Table S1:**
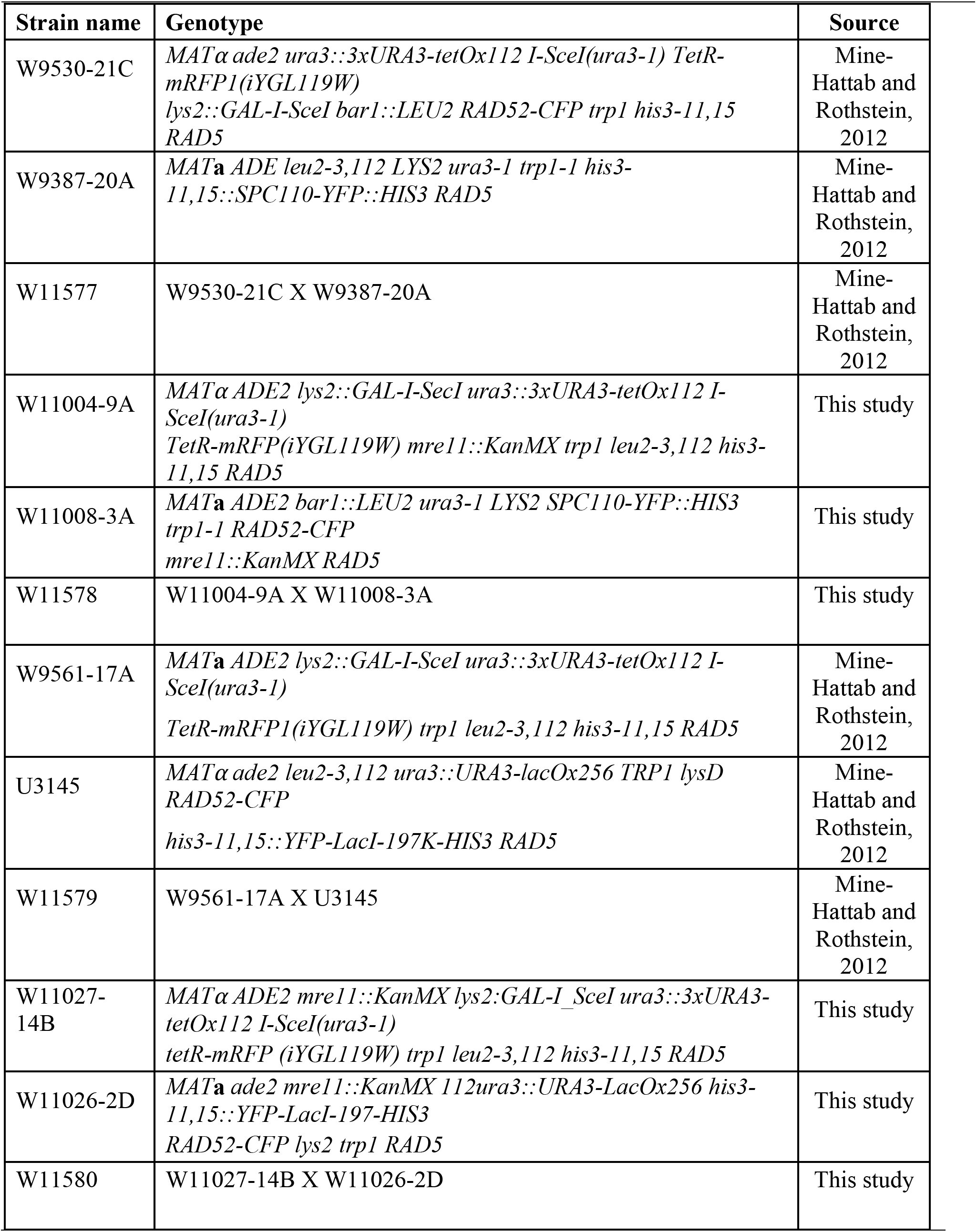
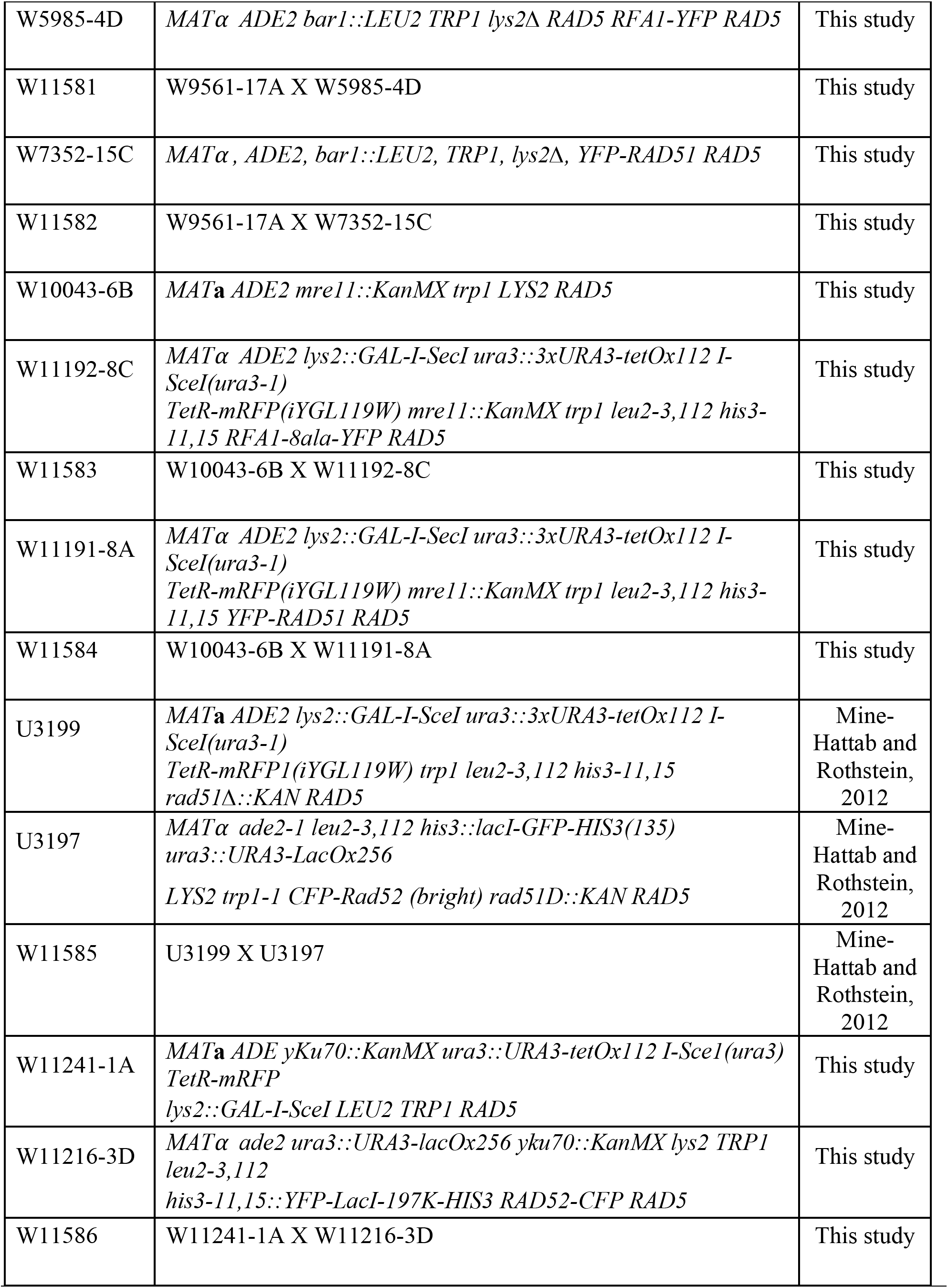
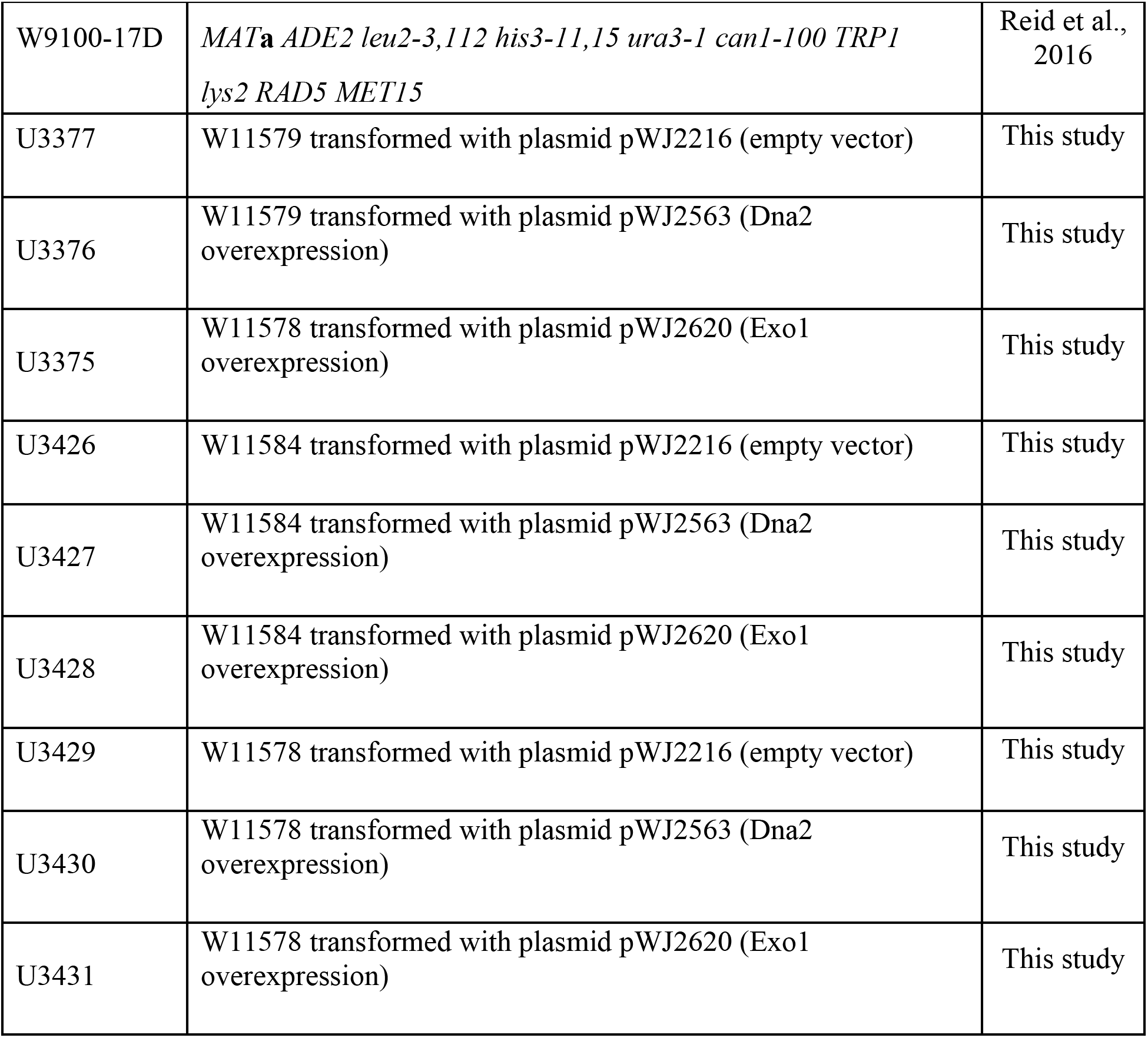
Yeast strains used in this study

**Table S2:**
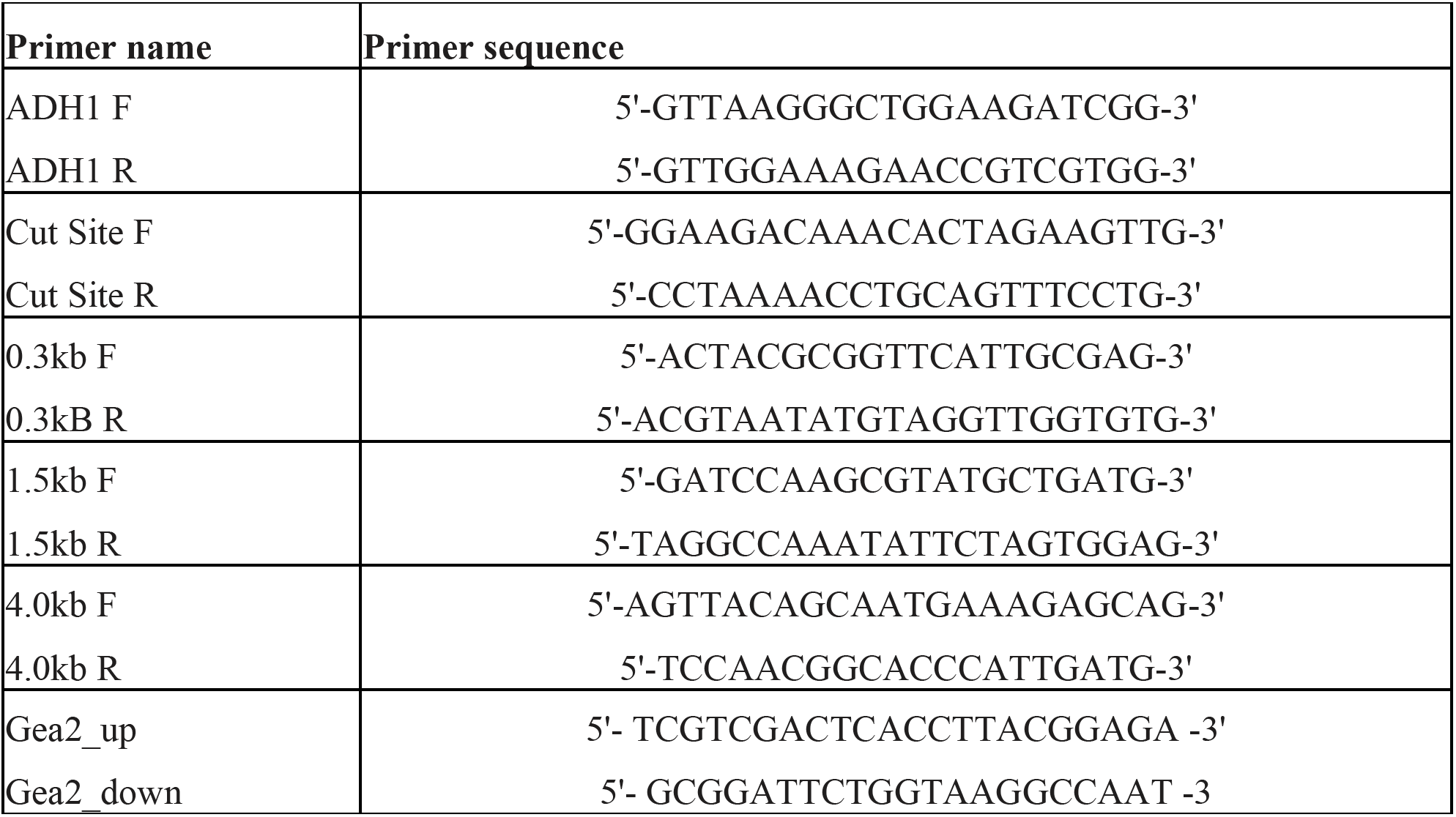
Primers and plasmids

**Table S3:** Summary of results

| Genotype | Experiment | Time after DSB induction | % measured |
| --- | --- | --- | --- |
| WT | I-SceI cutting | 0 | 0 |
|  |  | 30 | 22 |
|  |  | 60 | 55 |
|  |  | 90 | 56 |
|  |  | 120 | 54 |
|  |  | 150 | 48 |
|  |  | 180 | 38 |
|  |  | 240 | 23 |
|  |  | 300 | 20 |
|  |  | 360 | 15 |
|  |  | 420 | 4 |
| WT | Rfa1-foci | 0 | 0 |
|  |  | 90 | 24 |
|  |  | 120 | 35 |
|  |  | 150 | 46 |
|  |  | 180 | 61 |
|  |  | 240 | 59 |
|  |  | 300 | 59 |
| WT | Rad52-foci | 0 | 0 |
|  |  | 90 | 16 |
|  |  | 120 | 27 |
|  |  | 150 | 29 |
|  |  | 180 | 35 |
|  |  | 240 | 32 |
|  |  | 300 | 24 |
| WT | Rad51-foci | 0 | 0 |
|  |  | 90 | 9 |
|  |  | 120 | 33 |
|  |  | 150 | 40 |
|  |  | 180 | 44 |
|  |  | 240 | 54 |
|  |  | 300 | 53 |
| WT | Mobility | 0 | Confined |
|  |  | 120 | Linear |
|  |  | 180 | Linear |
| WT | Homolog Pairing | 0 | 2 |
|  |  | 90 | 4 |
|  |  | 120 | 19 |
|  |  | 150 | 21 |
|  |  | 180 | 17 |
|  |  | 240 | 13 |
|  |  | 300 | 7 |
| WT | Repair (GC) | 0 | 0 |
|  |  | 30 | 0 |
|  |  | 60 | 0 |
|  |  | 90 | 2 |
|  |  | 120 | 15 |
|  |  | 150 | 32 |
|  |  | 180 | 45 |
|  |  | 240 | 48 |
|  |  | 300 | 55 |
|  |  | 360 | 45 |
|  |  | 420 | 50 |

| Genotype | Experiment | Time of DSB induction | % measured |
| --- | --- | --- | --- |
| *mre11Δ* | I-SceI cutting | 0 | 0 |
|  |  | 30 | 32 |
|  |  | 60 | 65 |
|  |  | 90 | 75 |
|  |  | 120 | 79 |
|  |  | 150 | 77 |
|  |  | 180 | 80 |
|  |  | 240 | 64 |
|  |  | 300 | 47 |
|  |  | 360 | 40 |
|  |  | 420 | 35 |
| *mre11Δ* | Rfa1-foci | 0 | 0 |
|  |  | 90 | 3 |
|  |  | 120 | 5 |
|  |  | 150 | 11 |
|  |  | 180 | 9 |
|  |  | 240 | 27 |
|  |  | 300 | 34 |
| *mre11Δ* | Rad52-foci | 0 | 0 |
|  |  | 90 | 7 |
|  |  | 120 | 10 |
|  |  | 150 | 10 |
|  |  | 180 | 20 |
|  |  | 240 | 28 |
|  |  | 300 | 30 |
| *mre11Δ* | Rad51-foci | 0 | 0 |
|  |  | 90 | 2 |
|  |  | 120 | 5 |
|  |  | 150 | 6 |
|  |  | 180 | 8 |
|  |  | 240 | 38 |
|  |  | 300 | 41 |
| *mre11Δ* | Mobility | 0 | Confined |
|  |  | 180 | Confined |
|  |  | 240 | Linear |
| *mre11Δ* | Homolog Pairing | 0 | 3 |
|  |  | 90 | 3 |
|  |  | 120 | 5 |
|  |  | 150 | 2 |
|  |  | 180 | 6 |
|  |  | 240 | 15 |
|  |  | 300 | 20 |
| *mre11Δ* | Repair (GC) | 0 | 0 |
|  |  | 30 | 0 |
|  |  | 60 | 0 |
|  |  | 90 | 0 |
|  |  | 120 | 0 |
|  |  | 150 | 0 |
|  |  | 180 | 5 |
|  |  | 240 | 20 |
|  |  | 300 | 40 |
|  |  | 360 | 50 |
|  |  | 420 | 54 |
| *mre11Δ* +<br>Empty<br>vector | I-SceI cutting | 0 | 0 |
|  |  | 90 | 65 |
|  |  | 120 | 62 |
|  |  | 150 | 75 |
|  |  | 180 | 70 |
|  |  | 240 | 50 |
|  |  | 300 | 40 |
| *mre11Δ* +<br>Empty<br>vector | Rad51-foci | 0 | 0 |
|  |  | 90 | 0 |
|  |  | 120 | 0 |
|  |  | 150 | 0 |
|  |  | 180 | 10 |
|  |  | 240 | 38 |
|  |  | 300 | 41 |
| *mre11Δ* +<br>Empty<br>vector | Mobility | 0 | Confined |
|  |  | 180 | Confined |
| *mre11Δ* +<br>Empty<br>vector | Homolog<br>Pairing | 0 | 0 |
|  |  | 90 | 0 |
|  |  | 120 | 0 |
|  |  | 150 | 0 |
|  |  | 180 | 3 |
|  |  | 240 | 10 |
|  |  | 300 | 16 |
| *mre11Δ* +<br>Empty<br>vector | Repair (GC) | 0 | 0 |
|  |  | 90 | 0 |
|  |  | 120 | 0 |
|  |  | 150 | 0 |
|  |  | 180 | 5 |
|  |  | 240 | 22 |
|  |  | 300 | 35 |
| Genotype | Experiment | Time of DSB induction | % measured |
| *mre11Δ* +<br>Dna2 | I-SceI cutting | 0 | 0 |
|  |  | 90 | 65 |
|  |  | 120 | 62 |
|  |  | 150 | 75 |
|  |  | 180 | 70 |
|  |  | 240 | 50 |
|  |  | 300 | 40 |
| *mre11Δ + Dna2* | Rad51-foci | 0 | 0 |
|  |  | 90 | 0 |
|  |  | 120 | 14 |
|  |  | 150 | 24 |
|  |  | 180 | 42 |
|  |  | 240 | 53 |
|  |  | 300 | 59 |
| *mre11Δ + Dna2* | Mobility | 0 | Confined |
|  |  | 180 | Linear |
| *mre11Δ + Dna2* | Homolog Pairing | 0 | 0 |
|  |  | 90 | 0 |
|  |  | 120 | 6 |
|  |  | 150 | 8 |
|  |  | 180 | 15 |
|  |  | 240 | 10 |
|  |  | 300 | 5 |
| *mre11Δ + Dna2* | Repair (GC) | 0 | 0 |
|  |  | 90 | 0 |
|  |  | 120 | 0 |
|  |  | 150 | 15 |
|  |  | 180 | 25 |
|  |  | 240 | 40 |
|  |  | 300 | 42 |
| *mre11Δ + Exo1* | I-SceI cutting | 0 | 0 |
|  |  | 90 | 61 |
|  |  | 120 | 65 |
|  |  | 150 | 64 |
|  |  | 180 | 62 |
|  |  | 240 | 50 |
|  |  | 300 | 38 |
| *mre11Δ + Exo1* | Rad51-foci | 0 | 0 |
|  |  | 90 | 6 |
|  |  | 120 | 11 |
|  |  | 150 | 12 |
|  |  | 180 | 15 |
|  |  | 240 | 15 |
|  |  | 300 | 27 |
| *mre11Δ* +<br>Exo1 | Mobility | 0 | Confined |
|  |  | 180 | Confined |
| *mre11Δ* +<br>Exo1 | Homolog<br>Pairing | 0 | 0 |
|  |  | 90 | 0 |
|  |  | 120 | 0 |
|  |  | 150 | 0 |
|  |  | 180 | 0 |
|  |  | 240 | 16 |
|  |  | 300 | 4 |
| *mre11Δ* +<br>Exo1 | Repair (GC) | 0 | 0 |
|  |  | 90 | 0 |
|  |  | 120 | 0 |
|  |  | 150 | 0 |
|  |  | 180 | 5 |
|  |  | 240 | 27 |
|  |  | 300 | 40 |

For homolog pairing 5% background was removed for the summary plot in Figure 6.

## REFERENCES

Adkins, N. L., Niu, H., Sung, P., & Peterson, C. L. (2013). Nucleosome dynamics regulates DNA processing. Nat Struct Mol Biol, 20(7), 836–842. doi:10.1038/nsmb.2585

Brill, S. J., & Stillman, B. (1989). Yeast replication factor-A functions in the unwinding of the SV40 origin of DNA replication. Nature, 342(6245), 92–95. doi:10.1038/342092a0

Campbell, R. E., Tour, O., Palmer, A. E., Steinbach, P. A., Baird, G. S., Zacharias, D. A., & Tsien, R. Y. (2002). A monomeric red fluorescent protein. Proc Natl Acad Sci U S A, 99(12), 7877–7882. doi:10.1073/pnas.082243699

Cejka, P. (2015). DNA End Resection: Nucleases Team Up with the Right Partners to Initiate Homologous Recombination. J Biol Chem, 290(38), 22931–22938. doi:10.1074/jbc.R115.675942

Challa, K., Schmid, C. D., Kitagawa, S., Cheblal, A., Iesmantavicius, V., Seeber, A., Amitai, A., Seebacher, J., Hauer, M. H., Shimada, K., & Gasser, S. M. (2021). Damage-induced chromatome dynamics link Ubiquitin ligase and proteasome recruitment to histone loss and efficient DNA repair. Mol Cell, 81(4), 811–829 e816. doi:10.1016/j.molcel.2020.12.021

Cheblal, A., Challa, K., Seeber, A., Shimada, K., Yoshida, H., Ferreira, H. C., Amitai, A., & Gasser, S. M. (2020). DNA Damage-Induced Nucleosome Depletion Enhances Homology Search Independently of Local Break Movement. Mol Cell, 80(2), 311–326 e314. doi:10.1016/j.molcel.2020.09.002

Ciccia, A., & Elledge, S. J. (2010). The DNA damage response: making it safe to play with knives. Mol Cell, 40(2), 179–204. doi:10.1016/j.molcel.2010.09.019

Dion, V., & Gasser, S. M. (2013). Chromatin movement in the maintenance of genome stability. Cell, 152(6), 1355–1364. doi:10.1016/j.cell.2013.02.010

Dion, V., Kalck, V., Horigome, C., Towbin, B. D., & Gasser, S. M. (2012). Increased mobility of double-strand breaks requires Mec1, Rad9 and the homologous recombination machinery. Nat Cell Biol, 14(5), 502–509. doi:10.1038/ncb2465

Eapen, V. V., Sugawara, N., Tsabar, M., Wu, W. H., & Haber, J. E. (2012). The Saccharomyces cerevisiae chromatin remodeler Fun30 regulates DNA end resection and checkpoint deactivation. Mol Cell Biol, 32(22), 4727–4740. doi:10.1128/MCB.00566-12

Garcia Fernandez, F., Lemos, B., Khalil, Y., Batrin, R., Haber, J. E., & Fabre, E. (2021). Modified chromosome structure caused by phosphomimetic H2A modulates the DNA damage response by increasing chromatin mobility in yeast. J Cell Sci, 134(6). doi:10.1242/jcs.258500

Gasser, S. M. (2002). Visualizing chromatin dynamics in interphase nuclei. Science, 296(5572), 1412–1416. doi:10.1126/science.1067703

Gnügge, R., Oh, J., & Symington, L. S. (2018). Processing of DNA Double-Strand Breaks in Yeast. Methods Enzymol, 600, 1–24. doi:10.1016/bs.mie.2017.11.007

Goldstein, A. L., & McCusker, J. H. (1999). Three new dominant drug resistance cassettes for gene disruption in Saccharomyces cerevisiae. Yeast, 15(14), 1541–1553. doi:10.1002/(SICI)1097-0061(199910)15:14<1541::AID-YEA476>3.0.CO;2-K

Hauer, M. H., Seeber, A., Singh, V., Thierry, R., Sack, R., Amitai, A., Kryzhanovska, M., Eglinger, J., Holcman, D., Owen-Hughes, T., & Gasser, S. M. (2017). Histone degradation in response to DNA damage enhances chromatin dynamics and recombination rates. Nat Struct Mol Biol, 24(2), 99–107. doi:10.1038/nsmb.3347

Heim, R., & Tsien, R. Y. (1996). Engineering green fluorescent protein for improved brightness, longer wavelengths and fluorescence resonance energy transfer. Curr Biol, 6(2), 178–182. doi:10.1016/s0960-9822(02)00450-5

Herbert, S., Brion, A., Arbona, J. M., Lelek, M., Veillet, A., Lelandais, B., Parmar, J., Fernandez, F. G., Almayrac, E., Khalil, Y., Birgy, E., Fabre, E., & Zimmer, C. (2017). Chromatin stiffening underlies enhanced locus mobility after DNA damage in budding yeast. EMBO J, 36(17), 2595–2608. doi:10.15252/embj.201695842

Ivanov, E. L., Sugawara, N., White, C. I., Fabre, F., & Haber, J. E. (1994). Mutations in XRS2 and RAD50 delay but do not prevent mating-type switching in Saccharomyces cerevisiae. Mol Cell Biol, 14(5), 3414–3425. doi:10.1128/mcb.14.5.3414

Joseph, F., Lee, S. J., Bryant, E. E., & Rothstein, R. (2021). Measuring Chromosome Pairing During Homologous Recombination in Yeast. Methods Mol Biol, 2153, 253–265. doi:10.1007/978-1-0716-0644-5_18

Lamarche, B. J., Orazio, N. I., & Weitzman, M. D. (2010). The MRN complex in double-strand break repair and telomere maintenance. FEBS Lett, 584(17), 3682–3695. doi:10.1016/j.febslet.2010.07.029

Laurini, E., Marson, D., Fermeglia, A., Aulic, S., Fermeglia, M., & Pricl, S. (2020). Role of Rad51 and DNA repair in cancer: A molecular perspective. Pharmacol Ther, 208, 107492. doi:10.1016/j.pharmthera.2020.107492

Lawrimore, J., Barry, T. M., Barry, R. M., York, A. C., Friedman, B., Cook, D. M., Akialis, K., Tyler, J., Vasquez, P., Yeh, E., & Bloom, K. (2017). Microtubule dynamics drive enhanced chromatin motion and mobilize telomeres in response to DNA damage. Mol Biol Cell, 28(12), 1701–1711. doi:10.1091/mbc.E16-12-0846

Lee, C. S., Wang, R. W., Chang, H. H., Capurso, D., Segal, M. R., & Haber, J. E. (2016). Chromosome position determines the success of double-strand break repair. Proc Natl Acad Sci U S A, 113(2), E146–154. doi:10.1073/pnas.1523660113

Lee, S. E., Bressan, D. A., Petrini, J. H., & Haber, J. E. (2002). Complementation between N-terminal Saccharomyces cerevisiae mre11 alleles in DNA repair and telomere length maintenance. DNA Repair (Amst), 1(1), 27–40. doi:10.1016/s1568-7864(01)00003-9

Lewis, L. K., Karthikeyan, G., Westmoreland, J. W., & Resnick, M. A. (2002). Differential suppression of DNA repair deficiencies of Yeast rad50, mre11 and xrs2 mutants by EXO1 and TLC1 (the RNA component of telomerase). Genetics, 160(1), 49–62. Retrieved from https://www.ncbi.nlm.nih.gov/pubmed/11805044

Lisby, M., Barlow, J. H., Burgess, R. C., & Rothstein, R. (2004). Choreography of the DNA damage response: spatiotemporal relationships among checkpoint and repair proteins. Cell, 118(6), 699–713. doi:10.1016/j.cell.2004.08.015

Lisby, M., & Rothstein, R. (2009). Choreography of recombination proteins during the DNA damage response. DNA Repair (Amst), 8(9), 1068–1076. doi:10.1016/j.dnarep.2009.04.007

Lisby, M., & Rothstein, R. (2015). Cell biology of mitotic recombination. Cold Spring Harb Perspect Biol, 7(3), a016535. doi:10.1101/cshperspect.a016535

Marshall, W. F., Straight, A., Marko, J. F., Swedlow, J., Dernburg, A., Belmont, A., Murray, A. W., Agard, D. A., & Sedat, J. W. (1997). Interphase chromosomes undergo constrained diffusional motion in living cells. Curr Biol, 7(12), 930–939. doi:10.1016/s0960-9822(06)00412-x

Merigliano, C., & Chiolo, I. (2021). Multi-scale dynamics of heterochromatin repair. Curr Opin Genet Dev, 71, 206–215. doi:10.1016/j.gde.2021.09.007

Mimitou, E. P., & Symington, L. S. (2010). Ku prevents Exo1 and Sgs1-dependent resection of DNA ends in the absence of a functional MRX complex or Sae2. EMBO J, 29(19), 3358–3369. doi:10.1038/emboj.2010.193

Mine-Hattab, J., & Chiolo, I. (2020). Complex Chromatin Motions for DNA Repair. Front Genet, 11, 800. doi:10.3389/fgene.2020.00800

Mine-Hattab, J., Recamier, V., Izeddin, I., Rothstein, R., & Darzacq, X. (2017). Multi-scale tracking reveals scale-dependent chromatin dynamics after DNA damage. Mol Biol Cell. doi:10.1091/mbc.E17-05-0317

Mine-Hattab, J., & Rothstein, R. (2012). Increased chromosome mobility facilitates homology search during recombination. Nat Cell Biol, 14(5), 510–517. doi:10.1038/ncb2472

Mine-Hattab, J., & Rothstein, R. (2013). DNA in motion during double-strand break repair. Trends Cell Biol, 23(11), 529–536. doi:10.1016/j.tcb.2013.05.006

Moreau, S., Morgan, E. A., & Symington, L. S. (2001). Overlapping functions of the Saccharomyces cerevisiae Mre11, Exo1 and Rad27 nucleases in DNA metabolism. Genetics, 159(4), 1423–1433. doi:10.1093/genetics/159.4.1423

Niu, H., Chung, W. H., Zhu, Z., Kwon, Y., Zhao, W., Chi, P., Prakash, R., Seong, C., Liu, D., Lu, L., Ira, G., & Sung, P. (2010). Mechanism of the ATP-dependent DNA end-resection machinery from Saccharomyces cerevisiae. Nature, 467(7311), 108–111. doi:10.1038/nature09318

Ormo, M., Cubitt, A. B., Kallio, K., Gross, L. A., Tsien, R. Y., & Remington, S. J. (1996). Crystal structure of the Aequorea victoria green fluorescent protein. Science, 273(5280), 1392–1395. doi:10.1126/science.273.5280.1392

Reid, R. J., Du, X., Sunjevaric, I., Rayannavar, V., Dittmar, J., Bryant, E., Maurer, M., & Rothstein, R. (2016). A Synthetic Dosage Lethal Genetic Interaction Between CKS1B and PLK1 Is Conserved in Yeast and Human Cancer Cells. Genetics, 204(2), 807–819. doi:10.1534/genetics.116.190231

Rossetto, D., Avvakumov, N., & Cote, J. (2012). Histone phosphorylation: a chromatin modification involved in diverse nuclear events. Epigenetics, 7(10), 1098–1108. doi:10.4161/epi.21975

Schrank, B. R., Aparicio, T., Li, Y., Chang, W., Chait, B. T., Gundersen, G. G., Gottesman, M. E., & Gautier, J. (2018). Nuclear ARP2/3 drives DNA break clustering for homology-directed repair. Nature, 559(7712), 61–66. doi:10.1038/s41586-018-0237-5

Seeber, A., Dion, V., & Gasser, S. M. (2013). Checkpoint kinases and the INO80 nucleosome remodeling complex enhance global chromatin mobility in response to DNA damage. Genes Dev, 27(18), 1999–2008. doi:10.1101/gad.222992.113

Seeber, A., & Gasser, S. M. (2017). Chromatin organization and dynamics in double-strand break repair. Curr Opin Genet Dev, 43, 9–16. doi:10.1016/j.gde.2016.10.005

Sherman, F., Fink, G., & Hicks, J. B. (1987). Methods in Yeast Genetics: A Laboratory Course Manual: Cold Spring Harbor Laboratory Press.

Shim, E. Y., Chung, W. H., Nicolette, M. L., Zhang, Y., Davis, M., Zhu, Z., Paull, T. T., Ira, G., & Lee, S. E. (2010). Saccharomyces cerevisiae Mre11/Rad50/Xrs2 and Ku proteins regulate association of Exo1 and Dna2 with DNA breaks. EMBO J, 29(19), 3370–3380. doi:10.1038/emboj.2010.219

Shroff, R., Arbel-Eden, A., Pilch, D., Ira, G., Bonner, W. M., Petrini, J. H., Haber, J. E., & Lichten, M. (2004). Distribution and dynamics of chromatin modification induced by a defined DNA double-strand break. Curr Biol, 14(19), 1703–1711. doi:10.1016/j.cub.2004.09.047

Smith, M. J., Bryant, E. E., & Rothstein, R. (2018). Increased chromosomal mobility after DNA damage is controlled by interactions between the recombination machinery and the checkpoint. Genes Dev, 32(17-18), 1242–1251. doi:10.1101/gad.317966.118

Smith, M. J., & Rothstein, R. (2017). Poetry in motion: Increased chromosomal mobility after DNA damage. DNA Repair (Amst), 56, 102–108. doi:10.1016/j.dnarep.2017.06.012

Strecker, J., Gupta, G. D., Zhang, W., Bashkurov, M., Landry, M. C., Pelletier, L., & Durocher, D. (2016). DNA damage signalling targets the kinetochore to promote chromatin mobility. Nat Cell Biol, 18(3), 281–290. doi:10.1038/ncb3308

Sung, P. (1994). Catalysis of ATP-dependent homologous DNA pairing and strand exchange by yeast RAD51 protein. Science, 265(5176), 1241–1243. doi:10.1126/science.8066464

Sung, P., & Robberson, D. L. (1995). DNA strand exchange mediated by a RAD51-ssDNA nucleoprotein filament with polarity opposite to that of RecA. Cell, 82(3), 453–461. doi:10.1016/0092-8674(95)90434-4

Symington, L. S. (2016). Mechanism and regulation of DNA end resection in eukaryotes. Crit Rev Biochem Mol Biol, 51(3), 195–212. doi:10.3109/10409238.2016.1172552

Symington, L. S., Rothstein, R., & Lisby, M. (2014). Mechanisms and regulation of mitotic recombination in Saccharomyces cerevisiae. Genetics, 198(3), 795–835. doi:10.1534/genetics.114.166140

Thomas, B. J., & Rothstein, R. (1989). Elevated recombination rates in transcriptionally active DNA. Cell, 56(4), 619–630. doi:10.1016/0092-8674(89)90584-9

Tomita, K., Matsuura, A., Caspari, T., Carr, A. M., Akamatsu, Y., Iwasaki, H., Mizuno, K., Ohta, K., Uritani, M., Ushimaru, T., Yoshinaga, K., & Ueno, M. (2003). Competition between the Rad50 complex and the Ku heterodimer reveals a role for Exo1 in processing double-strand breaks but not telomeres. Mol Cell Biol, 23(15), 5186–5197. doi:10.1128/MCB.23.15.5186-5197.2003

Tsubouchi, H., & Ogawa, H. (2000). Exo1 roles for repair of DNA double-strand breaks and meiotic crossing over in Saccharomyces cerevisiae. Mol Biol Cell, 11(7), 2221–2233. doi:10.1091/mbc.11.7.2221

Usui, T., Ohta, T., Oshiumi, H., Tomizawa, J., Ogawa, H., & Ogawa, T. (1998). Complex formation and functional versatility of Mre11 of budding yeast in recombination. Cell, 95(5), 705–716. doi:10.1016/s0092-8674(00)81640-2

van Attikum, H., Fritsch, O., & Gasser, S. M. (2007). Distinct roles for SWR1 and INO80 chromatin remodeling complexes at chromosomal double-strand breaks. EMBO J, 26(18), 4113–4125. doi:10.1038/sj.emboj.7601835

Wright, W. D., Shah, S. S., & Heyer, W. D. (2018). Homologous recombination and the repair of DNA double-strand breaks. J Biol Chem, 293(27), 10524–10535. doi:10.1074/jbc.TM118.000372

Xue, C., Wang, W., Crickard, J. B., Moevus, C. J., Kwon, Y., Sung, P., & Greene, E. C. (2019). Regulatory control of Sgs1 and Dna2 during eukaryotic DNA end resection. Proc Natl Acad Sci U S A, 116(13), 6091–6100. doi:10.1073/pnas.1819276116

Yoder, T. J., Pearson, C. G., Bloom, K., & Davis, T. N. (2003). The Saccharomyces cerevisiae spindle pole body is a dynamic structure. Mol Biol Cell, 14(8), 3494–3505. doi:10.1091/mbc.e02-10-0655

Zhao, X., Muller, E. G., & Rothstein, R. (1998). A suppressor of two essential checkpoint genes identifies a novel protein that negatively affects dNTP pools. Mol Cell, 2(3), 329–340. doi:10.1016/s1097-2765(00)80277-4

Zhu, Z., Chung, W. H., Shim, E. Y., Lee, S. E., & Ira, G. (2008). Sgs1 helicase and two nucleases Dna2 and Exo1 resect DNA double-strand break ends. Cell, 134(6), 981–994. doi:10.1016/j.cell.2008.08.037

Zierhut, C., & Diffley, J. F. (2008). Break dosage, cell cycle stage and DNA replication influence DNA double strand break response. EMBO J, 27(13), 1875–1885. doi:10.1038/emboj.2008.111

Zou, L., & Elledge, S. J. (2003). Sensing DNA damage through ATRIP recognition of RPA-ssDNA complexes. Science, 300(5625), 1542–1548. doi:10.1126/science.1083430

